# Recording gene expression order in DNA by CRISPR addition of retron barcodes

**DOI:** 10.1101/2021.08.11.455990

**Authors:** Santi Bhattarai-Kline, Sierra K. Lear, Chloe B. Fishman, Santiago C. Lopez, Elana R. Lockshin, Max G. Schubert, Jeff Nivala, George Church, Seth L. Shipman

**Author notes:** **CORRESPONDING AUTHOR** Correspondence to S.L.S.

## Abstract

Biological processes depend on the differential expression of genes over time, but methods to make physical recordings of these processes are limited. Here we report a molecular system for making time-ordered recordings of transcriptional events into living genomes. We do this via engineered RNA barcodes, based on prokaryotic retrons^1^, which are reverse-transcribed into DNA and integrated into the genome using the CRISPR-Cas system^2^. The unidirectional integration of barcodes by CRISPR integrases enables reconstruction of transcriptional event timing based on a physical record via simple, logical rules rather than relying on pre-trained classifiers or post-hoc inferential methods. For disambiguation in the field, we will refer to this system as a Retro-Cascorder.

## INTRODUCTION

DNA is the universal storage medium for cellular life. In recent years, an emerging field of biotechnology has begun repurposing DNA to store data that has no cellular function. The same qualities of DNA that are beneficial in a biological context – high information density, ease of copying, and durability – also enable flexible storage of text, images, and sound^3-6^. Extending this general concept, researchers have developed data storage systems contained within living organisms that allow the recording of biological signals into DNA, such as endogenous transcription and environmental stimuli. One particular avenue of interest for such systems is in the longitudinal recording of biological processes within cells^7-9^.

These recordings address a fundamental limitation in standard methods to interrogate complex biological processes that require the destruction of cells and, thus, can only provide measurements at single points in time (e.g. RNA-Seq). Because biological processes are not perfectly synchronized at the cellular level, any individual cell collected in the middle of a biological process could be either ahead or behind in the progression of events relative to any other cell collected at that same time. This cellular heterochronicity makes it impossible to definitively reconstruct time-dependent processes from the destructive measurement of parallel samples. Indeed, cell-to-cell heterochronicity has actually been exploited in computational methods to infer position in a biological process among cells within a single sample (e.g. single-cell RNA-Seq pseudotime)^10^. However, these methods of inference make assumptions about the relationship between cells that are not explicitly known, and often require user-imposed constraints or the incorporation of prior biological knowledge^11^.

An approach known as molecular recording provides an alternative to statistical inference. Molecular recorders are biological devices that continuously record cellular processes, storing a physical record of the data permanently in cellular DNA, so that it may be retrieved at the very end of an experiment or process. Approaches to build molecular recorders have relied on different methods of modifying DNA, including site-specific recombinases and CRISPR-Cas nucleases^7,12,13^. Another approach to molecular recording, which we have worked to develop, leverages CRISPR-Cas integrases^14^. CRISPR-Cas integrases have been previously used to encode information into CRISPR arrays through the delivery of chemically synthesized oligos^4,14^ or by modulating the copy number of a reporter plasmid in response to a biological stimulus^5,8^. However, the ability to record the temporal order of more than one different biological signals into the CRISPR array of a single cell has not yet been demonstrated.

Here, we demonstrate successful recording of temporal relationships by adding a new molecular component to the system: a retroelement called a retron. The compact size, specificity, and flexibility of retrons to produce customizable DNA *in vivo* make them an attractive tool for biotechnology. Previously, retrons have been used in applications such as genome editing in several host systems^15-18^ and early analog molecular recorders^19^. By combining the functions of retrons and CRISPR-Cas integrases, we have built a system to make temporal recordings of transcriptional events.

To record transcriptional events, we engineered retrons to produce a set of compact, specific molecular tags, which can be placed under the control of multiple promoters of interest inside a single cell. When a tagged promoter is active, the tag sequence is transcribed into RNA, and reverse transcribed by the retron RT to generate a DNA ‘receipt’ of transcription. That DNA ‘receipt’ is then bound by Cas1-Cas2 and integrated into the cell’s CRISPR array, creating a permanent record of transcription. If another tagged promoter subsequently becomes active, a different DNA ‘receipt’ can be generated and integrated into the CRISPR array following the first spacer. By producing a linear record of these ‘receipts’ in the genome, we have built a biological device, called a Retro-Cascorder, that records the temporal history of specific gene expression events into the CRISPR arrays of individual cells (Fig. 1a).

**Figure 1.**
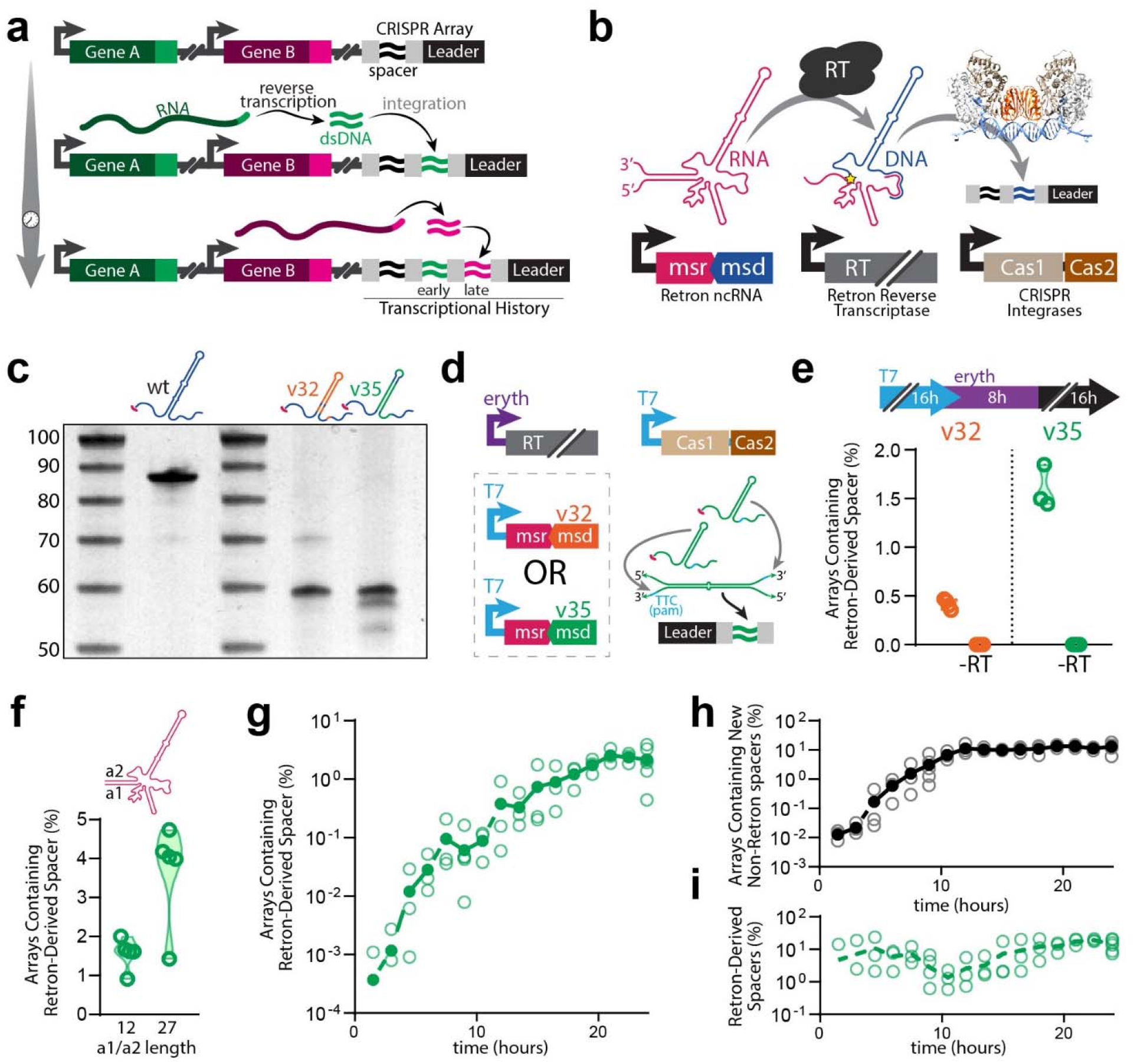
Cas1-Cas2 integrates retron RT-DNA. **a**. Schematic representation of retroelement-based transcriptional recording into CRISPR arrays. **b**. Schematic representation of biological components of the retron-based recorder. **c**. Urea-PAGE visualization of RT-DNA from retron Eco1 ncRNA variants. From left to right (excluding ladders): wild-type Eco1, Eco1 v32, Eco1 v35. For gel source data, see Supplementary Figure 1. **d**. Schematic of experimental promoters used to test retron-recorder parts and cartoon of hypothetical duplex RT-DNA prespacer structure. **e**. Quantification of arrays expanded with retron-derived spacers using Eco1 variants v32 (orange) and v35 (green). Open circles represent 3 biological replicates. **f**. Quantification of arrays expanded with retron derived spacers with a wild-type (12 bp) and extended (27 bp) a1/a2 region. Open circles represent 5 biological replicates. **g**. Time series of array expansions from retron-derived spacers. Open circles represent biological replicates, closed circles are the mean. **h**. Time series of array expansions from non-retron-derived spacers. Open circles represent biological replicates, closed circles are the mean. **i**. Proportion of total new spacers that are retron-derived. Open circles represent biological replicates, dashed line is the mean. All statistics in Supplementary Table 1.

## RESULTS

### Cas1-Cas2 integrates retron RT-DNA

CRISPR-Cas systems function as adaptive immune systems in bacteria and archaea. During the first phase of the immune response to infection by phage or mobile genetic elements, called adaptation, the CRISPR proteins Cas1 and Cas2 integrate a piece of foreign DNA into a genomic CRISPR array. The CRISPR array consists of a leader sequence followed by unique spacer sequences derived from foreign DNA, which are all separated by identical sequences called repeats. The sequence information stored in the spacers serves as an immunological memory of previous infection. This machinery, comprised of the CRISPR array, Cas1, and Cas2, is a ready-made storage device. When the Cas1-Cas2 complex integrates a spacer into the CRISPR array, it is added next to the leader sequence and the previous spacers are shifted away from the leader^20,21^. Thus, spacers which are further away from the leader sequence were acquired further in the past, and those closer to the leader acquired more recently.

The first challenge in building a temporal recorder of gene expression was to generate specific DNA barcodes following a transcriptional event, which can be permanently stored in a cell’s genome via integration by Cas1-Cas2. For integration, Cas1-Cas2 require DNA of at least 35 bases from end-to-end, with a 23 base complementary core region, and a protospacer-adjacent motif (PAM)^22^. To generate these acquirable pieces of DNA on-demand in cells, we used retrons. Recently determined to function in bacteria as a defense system against phage infection^23^, a typical retron consists of a single operon that controls the expression of: (1) a small, highly structured noncoding RNA (retron ncRNA), (2) a retron reverse transcriptase (RT) that specifically recognizes and reverse-transcribes part of its cognate ncRNA, and (3) one or more effector proteins which are implicated in downstream functions^23,24^. We designed variant ncRNA sequences of a native *E. coli* retron, Eco1^1,25^ (Ext. Data Fig. 1a), for integration into the genome by the type I-E CRISPR system of *E. coli* BL21-AI cells^20^ after they are reverse transcribed (Fig. 1b).

We tested multiple variants of Eco1 ncRNA for both reverse-transcription functionality and the ability of their RT-DNA to be acquired by the CRISPR adaptation machinery, and identified two that accomplish these aims. When overexpressed in *E. coli*, variants v32 and v35 (Ext. Data Fig. 1b-c) produced robust levels of RT-DNA that could be easily visualized on a PAGE gel (Fig. 1c), had perfect 3’-TTC PAM sequences, and could theoretically hybridize to create a 23-base core. Rather than a single copy of the retron RT-DNA hairpin forming the prespacer for acquisition, both v32 and v35 are designed such that two copies of the RT-DNA can form a duplex, which we hypothesized would be efficiently integrated into the CRISPR array (Fig. 1d, Ext. Data Fig. 1b-c). To measure the ability of variant retrons to be acquired, we overexpressed the variant ncRNA, Eco1 RT, and Cas1-Cas2 in BL21-AI cells which harbor a single CRISPR array in their genome. We then sequenced the CRISPR arrays of these cells to quantify integrations. In both cases, we found new spacers in these cells that matched the sequence of the retron RT-DNA that was expressed (Fig. 1e). Critically, arrays containing retron-derived spacers were only seen when cells also harbored a plasmid coding for Eco1 RT (Fig. 1e), indicating that the retron-derived spacers were indeed a result of the production of RT-DNA, rather than being derived exclusively from plasmid DNA. Retron v35 was acquired at a higher rate than v32 (Fig. 1e), and was selected for use in subsequent work.

We further modified v35 by extending the length of the non-hairpin duplex region referred to as the a1/a2 region (Fig. 1f). We have previously shown that this modification to retrons both increases production of RT-DNA in bacteria and yeast and increases the efficiency of genome editing methods which rely on retrons^18^. Consistent with our previous findings, extending the a1/a2 region of retron v35 resulted in an increase in the percentage of arrays which contained retron-derived spacers (Fig. 1f). This suggests that, like RT-DNA-templated genome editing, the rate of acquisition of retron-derived spacers is dependent on the abundance of RT-DNA. To take advantage of this improved acquisition efficiency, we incorporated this modification into all future Eco1 constructs.

To better characterize the acquisition of spacers by Cas1-Cas2 over time, we expressed retron v35 and Cas1-Cas2 for 24 hours and sampled arrays at regular intervals throughout (Fig. 1g-i). This showed that the number of arrays that contain retron-derived spacers increased regularly over time (Fig. 1g). As retron-derived spacers accumulated, they were accompanied by spacers derived from the cell’s genome and from plasmids, as previously described^20^. These non-retron-derived spacers also increased in arrays over time (Fig. 1h). The proportion of new retron-derived spacers remained relatively stable over time, making up between 1-10% of new spacer acquisitions (Fig. 1i). Thus, the abundance of retron-derived spacers can be used as a proxy for the duration of a transcriptional event. This result demonstrates a new implementation of analog molecular recording, similar in function to those previously described^19^, but based on the marriage of retrons and CRISPR-Cas integrases.

### Diversification of retron-based barcodes

A crucial advantage of retron-based molecular recording is the ability to follow multiple transcripts of interest by capturing distinct events within a single genomic CRISPR array. This enables the recording of gene expression timing within genetically identical cells, rather than relying on a mixed population of cells, each harboring different sensors. The specificity of retrons also enables more focused recordings compared to promiscuous RTs, which cannot be made to selectively reverse-transcribe individual transcripts^9,26^. Additionally, in contrast to recombinase-based molecular recording systems, the retron-based approach should enable a much larger set of sensors to coexist within a population of genetically identical cells. This is because the set of barcoded retrons is only limited by DNA sequence, rather than by the comparatively small number of well-characterized recombinases^7,27^. To construct a set of unique retron tags, we chose to use the loop in retron v35’s RT-DNA hairpin as a six-base barcode (Fig. 2a, Ext. Data Fig. 2). This barcoding strategy allows multiple otherwise identical ncRNAs to be reverse transcribed by the same RT, but remain easily distinguishable by sequence in CRISPR arrays. We synthesized a set of barcoded retrons, expressed them in cells along with Cas1-Cas2, and analyzed how efficiently they were acquired by sequencing CRISPR arrays (Fig. 2b). We compared these barcoded variants to the original v35 retron, and included a dead-RT version of the v35 retron as a negative control. Overall, we observed differences in the rate at which different barcoded retrons were acquired, ranging from no significant difference up to a ∼70% reduction in acquisitions (Fig. 2b). The differences in acquisition efficiency of the different barcoded retrons may come from changes in the efficiency of RT-DNA production, which we have observed to occur when changing the stem and loop region of retron ncRNAs^18^, and changes in the efficiency of acquisition, which we have observed to occur with different prespacer sequences^14^.

**Figure 2.**
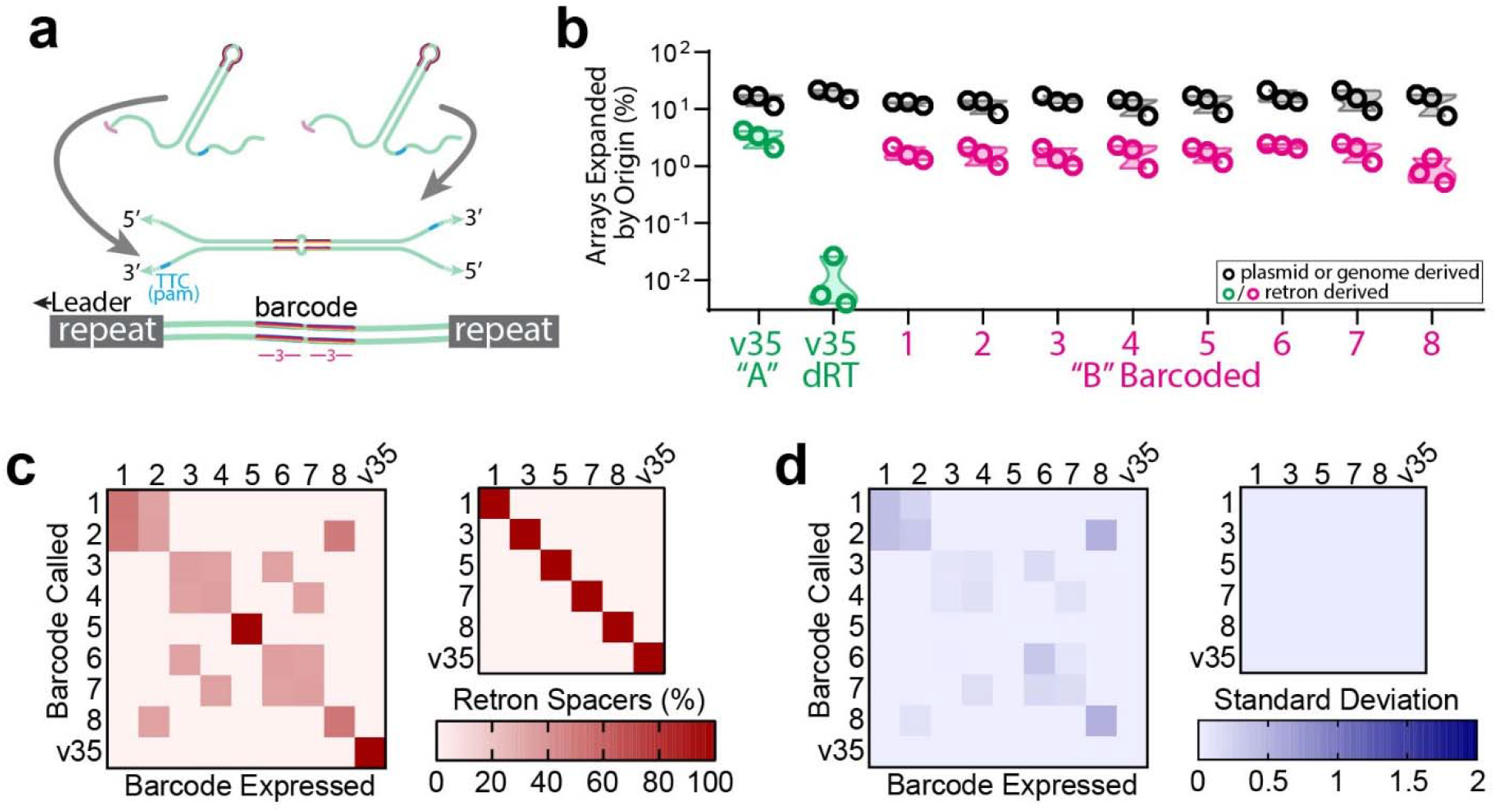
Diversification of retron-based barcodes. **a**. Hypothetical structure of duplexed RT-DNA prespacer with 6-base barcode and retron-derived spacer. **b**. Quantification of array expansions from barcoded variants of retron Eco1 v35, showing both retron-derived (green/pink) and non-retron derived (black) spacers for each variant. Open circles represent 3 biological replicates. **c**. Left: Heatmap of *in silico* ability to distinguish between all barcoded Eco1 v35 variants. Right: Heatmap of *in silico* ability to distinguish between reduced set of barcoded Eco1 v35 variants. **d**. Heatmap of standard deviation between three separate trials of barcode discrimination test. Left: full set. Right: reduced set. All statistics in Supplementary Table 1.

To test our ability to discriminate between the barcoded spacers derived from this set of retrons, we searched the sequence data from each sample expressing one barcoded retron for all of the other barcodes in the set. In our computational pipeline, we specify a tolerance of up to 3 bases of mismatches or indels (out of a 23 base search sequence) when determining the identity of a retron-derived spacer. This is to compensate for minor differences which may be found in mature spacers compared to their hypothetical sequence. As such, if our retron barcodes are faithfully preserved through all steps of the recording process (DNA coding sequence → RNA _→_ RT-DNA _→_ CRISPR array), then we should be able to effectively distinguish between barcodes which differ by 4 bases or more. This proved to be true when we examined our original set of 9 barcodes for orthogonality *in-silico*. Barcodes which differed by less than 4 bases could not be differentiated and barcodes which differed by 4 bases or more could be distinguished from each other with perfect accuracy, forming a set of 6 mutually orthogonal barcodes (Fig. 2c-d). This demonstrated that barcode sequences in retron-based transcriptional tags are faithfully preserved throughout the process of molecular recording, allowing for the facile construction of sets of mutually orthogonal tags.

### Mechanism of RT-DNA spacer acquisition

While it has been demonstrated that Cas1-Cas2 can integrate prespacers consisting of two complementary strands of DNA into the CRISPR array^14^, recent evidence suggests that Cas1-Cas2 are capable of binding ssDNA and may in fact bind the two strands of a prespacer separately, in a stepwise fashion^28^. To date, all experimentally characterized retrons have been shown to use the 2’-OH from a conserved guanosine to initiate reverse-transcription^1,24^, leaving a 2’-5’ RNA-DNA linkage. We hypothesized that this unique feature of RT-DNA prespacers might allow us to further interrogate the mechanism of prespacer loading and spacer acquisition by Cas1-Cas2.

Unlike many prespacers examined in prior work, which generally form perfect DNA duplexes, the duplex which we believe is formed by our retron has three characteristic regions where the prespacer should contain mismatches. The first of these regions is a stretch of five bases which, after integration, is located closest to the leader sequence. We will refer to this region as the leader-proximal, or LP, region (Fig. 3a). Next, there is a single base mismatch which falls near the middle of the mature spacer. We will refer to this region as the middle, or M, region (Fig. 3a). Finally, the last of the mismatched regions is found, in the mature spacer, in the five bases furthest from the leader sequence. We will refer to this as the leader-distal, or LD, region (Fig. 3a). We found that in retron-derived spacers, the sequence of these mismatched regions either corresponded to one strand of our hypothesized prespacer duplex or the other (Fig.. 3b). In this analysis, we will refer to the two strands of the hypothetical prespacer as the (+) and (-) strands. In Eco1-derived spacers, the sequence in the LP region overwhelmingly corresponded to the (-) strand. This (-) strand contains the PAM-proximal 3’-end, which determines directionality^14^ and has been shown to be integrated second in the spacer integration process^28,29^. This pattern of preserving the PAM-derived 3’-end sequence in the LP region was also seen when cells were electroporated with a synthetic oligonucleotide version of the retron RT-DNA (Fig. 3b).

**Figure 3.**
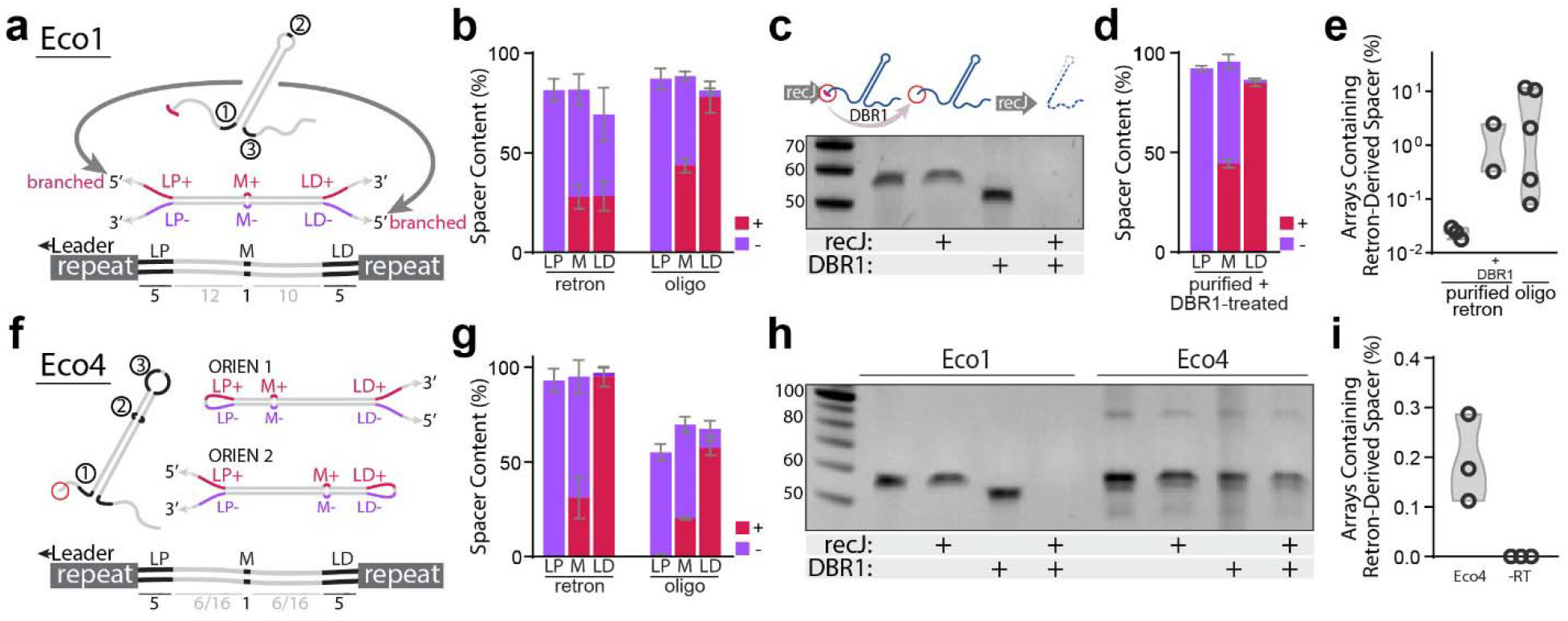
Mechanism of RT-DNA spacer acquisition. **a**. Hypothetical structure of duplexed Eco1 v35 RT-DNA prespacer and retron-derived spacer, with mismatched regions highlighted. **b**. Quantification of mismatch region sequences in spacers from cells expressing Eco1 v35 versus cells electroporated with oligo mimic. Bars represent the mean of 4 and 5 biological replicates for the retron and oligo-derived conditions, respectively (±SD). **c**. Urea-PAGE visualization of Eco1 RT-DNA. DBR1 treatment resolves 2’-5’ linkage. For gel source data, see Supplementary Figure 1. **d**. Quantification of mismatch region sequences in spacers from cells electroporated with purified, debranched Eco1 v35 RT-DNA. Bars represent the mean of 4 biological replicates (±SD). **e**. Quantification of array expansions from different prespacer substrates. Open circles represent 3, 2, and 5 biological replicates (left-right). **f**. Schematic of Eco4 RT-DNA, in both orientations, with mismatch sequences highlighted. **g**. Quantification of mismatch region sequences in cells expressing Eco4 versus cells electroporated with oligo mimic. Bars represent the mean of 3 biological replicates (±SD). **h**. Urea-PAGE visualization of Eco4 RT-DNA. DBR1 does not cause size shift of Eco4 RT-DNA. For gel source data, see Supplementary Figure 1. **i**. Quantification of array expansions from retron Eco4. Open circles represent 3 biological replicates (left-right). All statistics in Supplementary Table 1.

At the opposite end of the spacer, however, retron-derived and oligo-derived spacers were not identical. In the LD region, oligo-derived spacers overwhelmingly mapped to the (+) strand of our hypothesized duplex, whereas the LD regions of retron RT-DNA-derived spacers predominantly mapped to the (-) strand (Fig. 3b). Because the *in vivo*-produced RT-DNA contains a 2’-5’ linkage and the oligo does not, we suspected that the 2’-5’ linkage present in the Eco1 RT-DNA may interfere with the CRISPR adaptation process. To test this, we treated purified Eco1 RT-DNA with the eukaryotic debranching enzyme DBR1, which natively processes RNA lariats by cleaving 2’-5’ bonds in RNA^30^. Treatment of Eco1 RT-DNA with DBR1 *in vitro* resulted in a characteristic downward shift in the size of Eco1 RT-DNA from the loss of a small number of ribonucleotides remaining at the branch point. DBR1 treatment also rendered Eco1 RT-DNA sensitive to the 5’-exonuclease recJ (Fig. 3c). This indicates that DBR1 is able to remove the 2’-5’ linkage and produce Eco1 RT-DNA with an unbranched 5’-end. When purified Eco1 RT-DNA was treated with DBR1 and electroporated back into cells expressing Cas1-Cas2, the LD sequences of retron-derived spacers closely resembled those of retron-derived spacers after oligo electroporation (Fig. 3d), indicating that the presence of the 2’-5’ linkage in Eco1 RT-DNA is responsible for its unique pattern of spacer sequences. One potential explanation for the spacer pattern observed in Figure 3b is that, in addition to duplexed retron RT-DNA, the integrases may also bind and integrate prespacers consisting of one molecule of RT-DNA as the (-) strand and one molecule of plasmid-derived ssDNA (the retron coding sequence) as the (+) strand.

Beyond the apparent difference in prespacer processing due to the 2’-5’ linkage, we were curious to see whether the efficiency of acquisition would increase if the 2’-5’ linkage was removed. We approached this question by electroporating cells with three different prespacer types: purified RT-DNA, purified and debranched RT-DNA, and a synthetic oligo version of the RT-DNA. Debranched RT-DNA and oligos tended to be acquired more efficiently than the natively-branched RT-DNA, but this trend did not reach statistical significance (Fig. 3e).

In some retrons, processing naturally occurs following reverse transcription to remove the 2’-5’ linkage^31^, so we tested such a retron to see whether this processing would change the pattern or efficiency of retron-derived acquisitions. While the biosynthesis of the retron Eco4 RT-DNA still depends on priming from the 2’-hydroxyl of a conserved guanosine, its RT-DNA is cleaved 4 bases away from the 5’ branch point by an ExoVII exonuclease complex-dependent mechanism^31,32^, leaving a mature RT-DNA lacking a 2’-5’ linkage (Ext. Data Fig. 3a)^31^. We expressed wildtype Eco4 ncRNA, Eco4 RT, and Cas1-Cas2 in cells and then sequenced their CRISPR arrays to measure acquisitions. Notably, unlike the variant Eco1 retron, acquisitions from Eco4 occurred in two different orientations (Fig. 3f, Ext. Data Fig. 3b-c). Although the wildtype Eco4 RT-DNA does not have any perfect PAM sites (3’-TTC), both orientations observed in Eco4-derived spacers had a near-perfect PAM (3’-GTC) which proved sufficient for integration. We found no evidence as to whether these Eco4-derived spacers were derived from single hairpins or duplexes. We next analyzed the mismatched regions of the Eco4-derived spacers. As expected, almost all the LP regions mapped to the (-) strand, but unlike with variant Eco1, the LD region of Eco4-derived spacers almost entirely mapped to the (+) strand (Fig. 3g). Oligo-derived Eco4 spacers produced similar patterns of acquisition (Fig. 3g), indicating that retron Eco4, and likely other unbranched RT-DNAs, avoid the peculiarities caused by using a branched RT-DNA as a prespacer.

To confirm that Eco4 RT-DNA is debranched *in vivo*, we treated purified Eco4 RT-DNA with DBR1 and did not observe a size shift that would indicate removal of ribonucleotides (Fig. 3h). In addition, the RT-DNA was not recJ sensitive because there were fewer than 6 bases of single stranded DNA on the 5’ end, which recJ requires for exonuclease activity^33^.

The final test for Eco4 was to determine the overall efficiency of acquisition. We observed that retron-derived spacers from Eco4 were dependent on the presence of Eco4 RT, but their frequency was ultimately lower than Eco1-derived spacers (Fig. 3i). Based on these baseline efficiencies, we have focused our efforts on engineering Eco1 for the purpose of molecular recording. However, these results demonstrate that other retrons can also be used for molecular recording and, as is the case with Eco4, may possess unique qualities which affect their function in these applications.

### Temporal recordings of gene expression

Having built and characterized the requisite tools, we set out to make a temporal recording of gene expression using retron-based tags. We first constructed a signal plasmid and a recording plasmid. The recording plasmid contained the coding sequence for retron Eco1 RT, expressed from the constitutive promoter J23115, and the coding sequences for Cas1 and Cas2, both under the control of a T7/lac promoter. The signal plasmid, pSBK.134, harbored two copies of the Eco1 v35 ncRNA with different barcodes in the loop, which we will refer to as “A” and “B”, under different inducible promoters. “A” was under the control of the anhydrotetracycline-inducible promoter, pTet*, and ncRNA “B” was under the control of the choline chloride-inducible promoter, pBetI (Fig. 4a)^34^. We tested both the pTet* and pBetI promoters individually using YFP fluorescence and confirmed that both are responsive to their respective inducers with a similar maximum fluorescence, although the pBetI is ‘leakier’ with higher uninduced fluorescence (Ext. Data Fig. 4). We found no effect on growth of harboring these plasmids, and no effect on growth of adding inducers to cells that harbor the plasmids, but a small effect of inducing the pTet* promoter versus cells that harbor no plasmids (Ext. Data Fig. 5). The recorded responses to induction of pTet* and pBetI were well matched, with 24 hours of induction of each promoter yielding similar numbers of “A” and “B” derived spacers (Fig. 4b).

**Figure 4.**
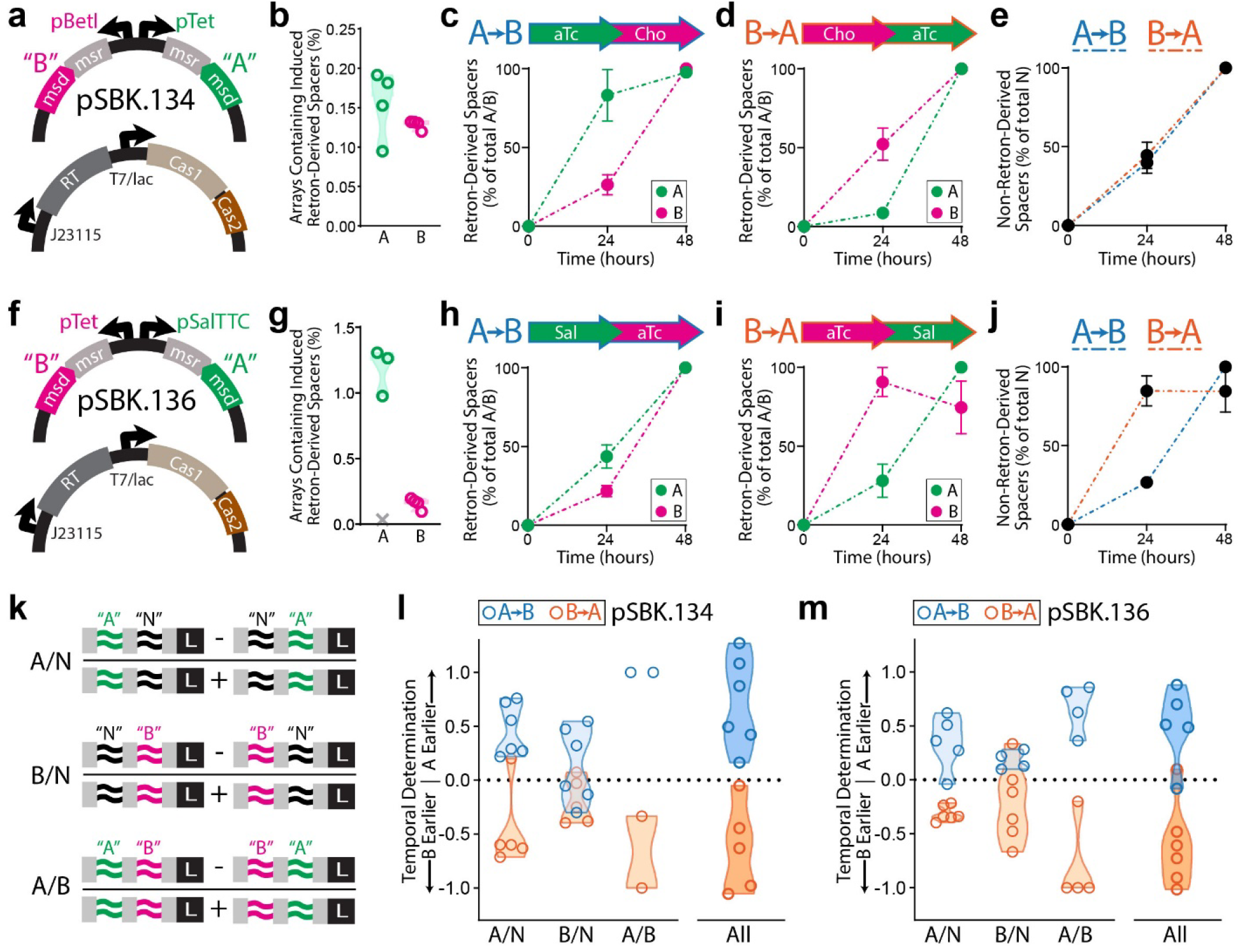
Temporal recordings of gene expression. **a**. Schematic of signal plasmid pSBK.134 used to express ncRNAs “A” and “B”, and recording plasmid used to express Eco1-RT and Cas1 and 2. **b**. Accumulation of retron-derived spacers from pSBK.134 after 24 hours of induction from their respective promoters (4 biological replicates). **c**. Retron-derived spacers when ncRNAs were induced in the order “A” then “B” from pSBK.134. Filled circles represent the mean of four biological replicates (±SEM). **d**. Retron-derived spacers when ncRNAs were induced in the order “B” then “A” from pSBK.134. Filled circles represent the mean of four biological replicates (±SEM). **e**. Non-retron-derived spacers in cells harboring pSBK.134, in both induction conditions. Filled circles represent the mean of four biological replicates (±SEM). **f**. Schematic of signal plasmid pSBK.136 used to express ncRNAs “A” and “B”, and the recording plasmid. **g**. Accumulation of retron-derived spacers from pSBK.136 after 24 hours of induction from their respective promoters (4 biological replicates). Outlier sample determined by Grubbs’ test denoted as a grey “X”. **h**. Retron-derived spacers when ncRNAs were induced in the order “A” then “B” from pSBK.136. Filled circles represent the mean of three biological replicates (±SEM). **i**. Retron-derived spacers when ncRNAs were induced in the order “B” then “A” from pSBK.136. Filled circles represent the mean of four biological replicates (±SEM). **j**. Non-retron-derived spacers in cells harboring pSBK.136, in both induction conditions. Filled circles represent the mean of four biological replicates (±SEM). **k**. Graphical representation of the rules used to determine order of expression from arrays. **l**. Ordering analysis of recording experiments with signal plasmid pSBK.134. Open circles are 6 biological replicates. **m**. Ordering analysis of recording experiments with signal plasmid pSBK.136. Open circles are 5 biological replicates. All statistics in Supplementary Table 1.

To record a time-ordered biological event, we transformed *E. coli* BL21-AI cells with both the signal and recording plasmids, and grew them under two different experimental conditions for a total of 48 hours. In the first temporal recording condition, cells were grown for 24 hours with inducers driving the expression of Eco1 RT, Cas1-2, and ncRNA “A”. The cells were then grown for another 24 hours while expressing ncRNA “B”, along with the Eco1 RT and Cas1-Cas2 (Fig. 4c). In the second condition, the order of expression of ncRNA “A” and “B” was reversed (ncRNA “B” was expressed for the first day and ncRNA “A” for the second) (Fig. 4d). Samples were taken at 24 and 48 hours. Examination of the expanded arrays revealed a significant increase in the percentage of cells that received a retron-derived spacer in the 24 hours where its chemical inducer was present, compared to the 24 hours where it was absent. This held true for both ncRNAs “A” and “B” under both the “A”-before-”B” and “B”-before-”A” expression schemes (Fig. 4c-d). The number of non-retron-derived spacers also increased consistently over 48 hours (Fig. 4e).

To further test the generalizability of the system, we made a recording of a different set of promoters driving the same retron ncRNAs. For this second arrangement, the recording plasmid remained the same, but in the signal plasmid, pSBK.136, ncRNA “A” was placed under the control of the sodium salicylate-inducible promoter, pSal, and “B” under the control of pTet* (Fig. 4f). We also validated the pSal promoter individually using YFP fluorescence and found it to be responsive to its inducer, with a higher maximum fluorescence than the pTet* promoter (Ext. Data Fig. 4). Consistent with this difference, 24 hours of induction of each promoter resulted in a much higher rate of acquisitions from retron “A” driven by pSal than acquisitions of retron “B” from pTet* (Fig. 4g). In this case, induction of the pSal promoter did result in a negative effect on population growth (Ext. Data Fig. 5). Notably, in one biological replicate, the recording system appeared to break, resulting in nearly non-existent acquisitions; this sample was excluded from further analysis following its identification as an outlier by Grubbs’ test^35,36^ (Fig. 4g). Next, we tested two experimental conditions: “A”-before-”B” and “B”-before-”A” (Fig. 4h-i). Despite the mismatched promoter strengths, when arrays were examined at the 24- and 48-hour timepoints, more arrays were expanded with retron-derived spacers in the presence of their respective inducers than in their absence (Fig. 4h-i). In addition, the numbers of non-retron-derived spacers again increased over 48 hours (Fig. 4j).

This analysis of spacer acquisitions from the signal plasmid was enabled by a timepoint sampling in the middle of the overall transcriptional sequence. However, the aim of this work is to reconstruct the timing of transcriptional events using only data acquired at an endpoint. Therefore, we defined logical rules that should govern the ordering of spacers in the CRISPR arrays, and allow us to reconstruct the order of transcription of separate ncRNAs. Because spacers are acquired unidirectionally, with newer spacers closer to the leader sequence, we postulated that if transcript “A” is expressed before transcript “B”, arrays of the form “A” _→_ “B” _→_ Leader should be more numerous than “B” _→_ “A” _→_ Leader. Accordingly, if “B” is expressed before “A”, then the opposite should be true: the number of “B” _→_ “A” _→_ Leader arrays should be greater than the number of “A” _→_ “B” _→_ Leader arrays.

Another feature of using CRISPR arrays for recording is that Cas1-2 also acquire spacers derived from the plasmid and genome^20^. These untargeted acquisitions can also be used to interpret temporal information^5,8^. If we assume that these non-retron-derived spacers (denoted “N”) are acquired at a constant rate throughout the experiment, we can define a set of rules that govern the order of “N” _→_ “A” _→_ Leader versus “A” _→_ “N” _→_ Leader arrays and of “N” _→_ “B” _→_ Leader versus “B” _→_ “N” _→_ Leader arrays. In the “A”-before-”B” case, since “A” is expressed in the first half of an experiment, arrays of the form “A” _→_ “N” _→_ Leader should be more numerous than “N” _→_ “A” _→_ Leader. And since “B” is expressed in the second half of the experiment, “N” _→_ “B” _→_ Leader arrays should be more numerous than “B” _→_ “N” _→_ Leader arrays. Likewise, in the “B”-before-”A” condition, “N” _→_ “A” _→_ Leader arrays should be more numerous than “A” _→_ “N” _→_ Leader arrays and “B” _→_ “N” _→_ Leader arrays should be more numerous than “N” _→_ “B” _→_ Leader arrays. Restating these as mathematical statements, we can take the difference between possible array types (e.g. “A” _→_ “B” _→_ Leader minus “B” _→_ “A” _→_ Leader) as the numerator and the sum of the two possibilities (e.g. “A” _→_ “B” _→_ Leader plus “B” _→_ “A” _→_ Leader) as the denominator (Fig. 4k) to yield a number between -1 and 1 for each ordering rule (A/B, A/N, and B/N). By the convention of our ordering rules, positive values would indicate that “A” was present before “B”, and a negative output would indicate that “B” was present before “A”. The magnitude of the output (0 _≤_ |x| _≤_ 1) is a measure of how strongly the rule is satisfied in a given direction, or in other words, how complete is the separation of the two signals in time.

To test these predictions, we sequenced the CRISPR arrays of all of our samples at the 48-hour endpoint. Across 6 biological replicates of samples with signal plasmid pSBK.134, the samples in which “A” was expressed before “B” yielded positive values when subjected to analysis by our ordering rules, correctly identifying the order of expression. Likewise, for samples where “B” was expressed before “A”, the rules yielded negative values, again correctly identifying the order (Fig. 4l). We also calculated a composite score by taking a weighted average of all three rules. This score consists of the average between the A/B rule and the sum of the A/N rule and B/N rule. We devised this formulation based on what the ordering rules represent in an ideal system. By definition, the A/B rule represents the degree of order between A and B and will have a magnitude between 0 and 1. When “A” and “B” are not at all ordered with respect to time the ordering score should be 0, and when “A” and “B” completely separated in time the ordering score should be 1. Likewise, the A/N and B/N rules represent the degree of order between “N” and “A” or “B”, respectively. In an ideal system, where the rate of acquisition of “N” is constant, the magnitude of the A/N and B/N scores should be constrained between 0 and 0.5, and the sum of the A/N score and B/N score can be used as a proxy for the order of “A” with respect to “B”. Thus, if we assume that the rate of acquisition of “N” is constant, we can average the sum of the A/N and B/N scores with the A/B score to generate a composite score which integrates all three rules and is representative of the degree of temporal order between A and B. It is important to note that in the *in vivo* recording data, there are samples in which the magnitudes of the A/N and/or B/N scores exceeds the ideal value of 0.5 and, as a result, the composite score exceeds 1. We believe that this could occur due to several reasons. One reason is that our assumption of “N” being a constant signal is not true *in vivo*, and that the strength of signal “N” has some structure in time. Another potential reason is that the recording of these signals is a stochastic process, with randomness and noise introduced at many levels of the system, from RT-DNA synthesis, to spacer acquisition, to cell division, to sampling.

When applied to our *in vivo* recording data, this method accurately determined that each experiment yielded directional acquisition of spacers and correctly recalled the order of events for both directions. Critically, this demonstrates our ability to accurately reconstruct the order of two transcriptional events in an endpoint biological sample, using only logical rules derived from first principles. Interestingly, the retron signal driven by the pBetI promoter, which was found to be leakier when uninduced, was not as strongly directional as the pTet*-driven signal in relation to N spacers, as would be expected. When each replicate was examined separately, though all rules were not uniformly satisfied, the order of expression could be consistently determined (Ext. Data Fig. 6a-b).

When this analysis was applied to samples with the signal plasmid pSBK.136 (which had mismatched “A” and “B” promoter strengths), we were still able to accurately reconstruct the order of events from endpoint data (Fig. 4m, Ext. Data Fig. 6c-d), demonstrating that the temporal analysis of gene expression can be generalized to different promoters.

Finally, Retro-Cascorder data is stably maintained in cells for multiple generations after the completion of a recording. When cells containing retron-derived recordings using signal plasmid pSBK.134 were passaged for multiple days, ordering analysis results remained very stable through roughly 18 generations of cell division (Ext. Data Fig. 7a-d). Only after around 45 generations of division did ordering scores begin to experience moderate drift, with severe drift apparent after 81 generations. For reference, the Hayflick limit of human fetal cells in vitro is 40 to 60 generations of cell division^37^. Ultimately, this paradigm enables the reconstruction of temporal histories within genetically-identical populations of cells, based on a physical molecular record.

### Modeling the Limits of Retron Recording

To better understand the nature of the retron recording system and its behavior in a wide range of conditions, including those which we are unable to recreate in the lab, we developed a computational model of the Retro-Cascorder based on data from our temporal recordings and present understanding of the biology of the system. Using the raw number of acquisitions observed previously from temporal recordings, at the 24- and 48-hour timepoints, we defined a set of rates (of acquisitions per hour) for the different signals recorded by the cells. Based on the overall low rate of acquisitions, we assume that the spacer acquisitions can be modeled faithfully as a Poisson process, wherein the average time between events (here acquisitions) is known but the exact timing of events is random. Using rates estimated from our recordings, we define rates of acquisition for A and B in the presence of their respective inducers, A and B in the absence of their respective inducers (to account for leak from their promoters), and a constant background rate of acquisition of non-retron-derived spacers, or N.

To test our model, we first simulated 100 replicates, of 1 million arrays each, of our previous recording experiments using the signal plasmids pSBK.134 and pSBK.136 (Fig. 5a-b). With the results of the simulation appearing to approximate the results from our real recordings, we next sought to understand how changing various parameters of the recording system may affect results. First, we simulated the effect of analyzing different numbers of arrays (Fig. 5c). The simulation suggests that it is important to dedicate a generous number of sequencing reads to a recording experiment in order to properly resolve the process in question. When too few arrays are analyzed from a given sample, the calculated ordering scores will be unreliable, as evidenced by the very wide distribution of composite scores from simulated low-read samples.

Next, we simulated making recordings which varied in length (Fig. 5d). The effect that appears here is similar to the effect of varying the number of arrays analyzed. In the range of very short recordings, the system is unable to resolve the order of the signals, but as the length of the recordings increases, the composite scores converge toward a specific value. To check this finding from the model, we made biological recordings of different length using pSBK.134. In these recordings, the trend predicted by the model appeared to hold, with shorter recordings unable to resolve the order of the signals and longer recordings with greater fidelity. Finally, we simulated the effect of varying the rates of acquisition of both retrons. To do this we simulated 50 replicates, of 1 million arrays each, across a range of rates of acquisition of both signals A and B. We varied both the induced and uninduced rates of a given signal by the same factor (e.g. A-On and A-Off increased by a factor of 4, B-On and B-Off decreased by a factor of 8). Interestingly, even when acquisition rates are decreased, the mean ordering scores across 50 replicates faithfully reflect the order of expression of signals (Fig. 5e-g). Dispersion of the ordering scores among replicates, however, varies dramatically with the rates of acquisition of signals A and B. In short: as the strength of signals A and B increases, we expect to be able to faithfully recall temporal order using fewer replicates, or visa-versa, that if the strength of signals decreases, more replicates will be required to resolve their temporal order.

**Figure 5.**
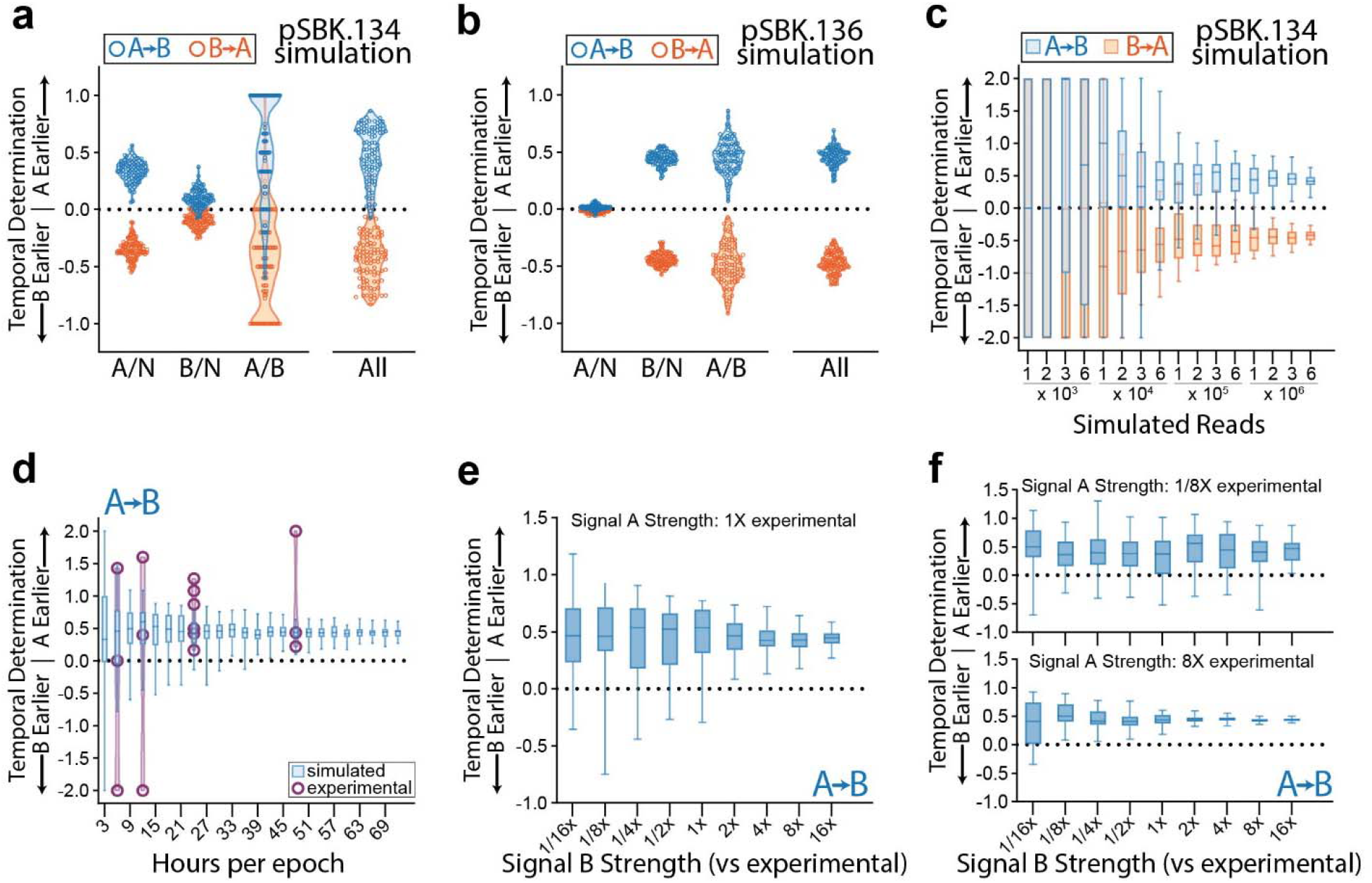
Modeling the Limits of Retron Recording. **a**. Simulation of 100 replicates each of A-then-B and B-then-A recordings using acquisition rate data from pSBK.134 recordings. Each point represents the calculated ordering score from a single replicate of 1 million arrays. **b**. Simulation of 100 replicates each of A-then-B and B-then-A recordings using acquisition rate data from pSBK.136 recordings. Each point represents the calculated ordering score from a single replicate of 1 million arrays. **c**. Simulation of varying the number of arrays analyzed per sample using acquisition rate data from pSBK.134 recordings. Each box with whiskers represents 100 simulated replicates, with whiskers extending from minimum to maximum. **d**. Simulation of varying the length of each epoch in a retron recording using acquisition rate data from pSBK.134 (blue). Overlaid with real retron recordings of the same length (purple). Each box with whiskers represents 100 simulated replicates of 1 million reads each, with whiskers spanning from minimum to maximum. Each overlaid point is a single biological replicate. Recording experiments with 6, 12, and 48-hour epochs were done in triplicate. Recording experiments with epoch length of 24 hours are the same as in Figure 4l. **e**. Simulation of varying the strength of signal B when signal A remains constant. 1x acquisition rates were obtained from pSBK.134 recordings. Each box with whiskers represents 50 simulated replicates of 1 million arrays each. Whiskers span from minimum to maximum. **f**. Simulation of varying the strength of signal B when signal A is decreased or increased by a factor of 8. 1x acquisition rates were obtained from pSBK.134 recordings. Each box with whiskers represents 50 simulated replicates of 1 million arrays each. Whiskers span from minimum to maximum.

Putting together the pieces above, we believe that we have shown 4 variables that are critical to the design of these recording experiments: (1) signal strength; (2) length of recording; (3) number of reads; and (4) number of replicates. By increasing any of these 4 parameters, the experimenter can expect greater fidelity of their final temporal recording. When these parameters are decreased, one should expect more noise and variability in their recordings. In the laboratory however, there will be practical limits as to how much the experimenter can maximize or alter these parameters. Often, it may not be possible to alter the duration of an experiment or the strength of a transcriptional signal due to the biology of the process of interest, and it may be time- and cost-prohibitive to run large numbers of biological replicates. Of the four parameters then, increasing the number of arrays analyzed (and consequently the number of sequencing reads) from individual samples is likely the cheapest and simplest way to increase the fidelity of temporal recordings and final analyses.

## DISCUSSION

Here, we have described a system for the recording and reconstruction of transcriptional history in a population of cells, which we call the Retro-Cascorder. We achieved this by engineering an RNA molecular tag, which is specifically reverse-transcribed to produce a DNA ‘receipt’ of transcription that is permanently saved in a CRISPR array. We demonstrated the flexibility and potential for continued development of these tools by making the recording retron more efficient with modifications to the structure of the retron ncRNA, and developed a toolkit of barcoded retrons for future application to more complex systems. Beyond this, we investigated the ability of the CRISPR adaptation system to utilize RT-DNA as a prespacer, and discovered that the retron 2’-5’ linkage causes a marked difference in the type of spacers acquired. Finally, we used this system to record and reconstruct time-ordered biological events in populations of cells, and developed a computational model of retron-recording to more comprehensively explore the limits of the system.

One natural aspect of the CRISPR integrases which has proven useful in these recordings is the acquisition of diverse spacers from plasmid and genomic fragments (N spacers). In our temporal recordings of two inducible elements, these N spacers function as a third signal, providing a constant background. Because the integrases are also driven by an inducible promoter, this background signal marks the timing of the recording components. In our recordings with pSBK.134, for instance, the A spacers encode the timing of anhydrotetracycline in the media, the B spacers encode the timing of choline chloride in the media, and the N spacer encode the timing of arabinose and IPTG in the media. The frequency of N spacer acquisitions is unaffected by the retron-derived acquisitions, which we interpret to mean that the acquisition of events in this system is not competitive, but rather additive. Therefore, these N spacers do not interfere with the recording, but rather aid in resolving the temporal order of recorded signals.

Here, we recorded two distinct signals within a homogenous population of cells, with the N spacers serving as a constant third signal. The level of complexity of these recordings is similar to previous work using different recombinases to encode events^7,27,38^. One aspect that is encouraging about the system described here is that the recordings use a common set of protein components, with distinct signals being encoded using variable nucleotides in the retron-derived spacers. Recombinase-based recorders, while robust, are inherently limited in the number of distinct signals to 2^number of recombinases^39^, which requires identifying and expressing many orthogonal recombinases. In contrast, this approach is limited in the number of distinct signals to 4^number of nucleotides used in the barcode. This bodes well for the scalability of this approach, but the practicality of scaling will need to be experimentally validated in future work.

We believe that this framework of selective tagging and recording of biological signals in an RNA _→_ DNA _→_ CRISPR direction is a powerful, modular, and extensible method of making temporal recordings in cells. Using only *a priori* ordering rules, we can detect and interpret time-ordered biological signals from a single endpoint sample. Immediate uses of this technology include the construction of living biosensors that sample and record their environment. Here, we record the presence of anhydrotetracycline, choline chloride, sodium salicylate, arabinose, and IPTG. Near future work could modify these systems to record the presence of pollutants, metabolites, or pathogens in an environment. With additional engineering to increase the efficiency of the recordings, we hope that this system will enable recordings of natural gene expression to log transcriptional order during complex cellular events.

## METHODS

All biological replicates were taken from distinct samples, not the same sample measured repeatedly.

### Bacterial Strains and Growth Conditions

This work uses the following *E. coli* strains: NEB 5-alpha (NEB C2987, not authenticated), BL21-AI (ThermoFisher C607003, not authenticated), bMS.346, and bSLS.114. bMS.346 was generated from *E. coli* MG1655 by inactivating *exoI* and *recJ* genes with early stop codons as in previous work^40^. Additionally, the *araB*::T7RNAP-*tetA* locus was transferred from BL21-AI by P1 phage transduction^41^. bSLS.114 (which has been used previously^18^) was generated from BL21-AI by deleting the retron Eco1 locus by lambda Red recombinase mediated insertion of an FRT-flanked chloramphenicol resistance cassette. This cassette was amplified from pKD3^42^ with homology arms added to the retron Eco1 locus. This amplicon was electroporated into BL21-AI cells expressing lambda Red genes from pKD46^42^, and clones were isolated by selection on chloramphenicol (10 μg/mL) plates. After genotyping to confirm locus-specific insertion, the chloramphenicol cassette was excised by transient expression of FLP recombinase to leave only an FRT scar. Experimental cultures were grown with shaking in LB broth at 37ºC with appropriate inducers and antibiotics. Inducers and antibiotics were used at the following working concentrations: 2 mg/mL L-arabinose (GoldBio A-300), 1 mM IPTG (GoldBio I2481C), 400 μM erythromycin, 100 ng/mL anhydrotetracycline, 100 μM choline chloride, 1 mM sodium salicylate, 35 μg/mL kanamycin (GoldBio K-120), 25 μg/mL spectinomycin (GoldBio S-140), 100 μg/mL carbenicillin (GoldBio C-103), 25 μg/mL chloramphenicol (GoldBio C-105; used at 10 μg/mL for selection during recombineering). Additional strain information can be found in Supplementary Table 2.

### Plasmid Construction

All cloning steps were performed in *E. coli* NEB 5-alpha. pWUR 1+2, containing Cas1 and Cas2 under the expression of a T7lac promoter, was a generous gift from Udi Qimron^20^. Eco1 wildtype ncRNA and Eco1 RT, along with Cas1+2, were cloned into pRSF-DUET (Sigma 71341) to generate pSLS.405. Eco1 variant ncRNA sequences v32 and v35 were cloned into pRSF-DUET along with Cas1+2 to generate pSLS.407 and pSLS.408, respectively. Extended a1/a2 v35 ncRNA expression plasmid pSLS.416 was generated from pSLS.408 by site-directed mutagenesis. Retron Eco1 RT and retron Eco4 RT were cloned into pJKR-O-mphR to generate pSLS.402 and pSLS.400, respectively. pJKR-O-mphR was generated previously^43^ (Addgene plasmid # 62570). Barcoded, extended a1/a2 v35 ncRNA expression plasmids pSBK.009-016 were generated from pSLS.416 by site-directed mutagenesis. Wildtype retron Eco4 ncRNA was cloned into pRSF-DUET along with Cas1+2 to generate SLS.419. pSBK.134 and pSBK.136 were generated in three steps. First, barcoded, extended a1/a2 v35 ncRNA sequences were cloned into the ‘Marionette’ plasmids pAJM.717, pAJM.718, and pAJM.771. pAJM.717, pAJM.718, and pAJM.771 were gifts from Christopher Voigt^34^ (pAJM.717 - Addgene plasmid # 108517 // pAJM.718 - Addgene plasmid # 108519 // pAMJ.771 - Addgene plasmid # 108534). Then, in two steps, two ncRNA expression cassettes (for barcoded ncRNAs “A” and “B”) from the Marionette plasmids were cloned into pSol-TSF (Lucigen F843213-1) facing in opposite directions. pSBK.079 was generated by cloning the resistance marker AmpR in place of the KanR marker into the plasmid pSLS.425, which was synthesized by Twist biosciences. Additional plasmid information can be found in Supplementary Table 3.

### RT-DNA Purification and Visualization

Retron RT-DNA was expressed in *E. coli* bMS.346 and purified in two steps. First, DNA was extracted from cells using a plasmid midiprep kit (Qiagen 12943). This purified DNA was then treated for 30 minutes at 37C with RNAse A/T1 mix (ThermoFisher EN0551) and, if required, DBR1 (OriGene TP300024) and/or RecJ_f_ (NEB M0264). This sample was then used as the input for the Zymo Research ssDNA/RNA Clean & Concentrate kit (Zymo D7011). Samples eluted from the ssDNA kit were resolved using TBE-urea PAGE (ThermoFisher EC6885BOX). Gels were stained with SYBR Gold for imaging (ThermoFisher S11494) and imaged on a Bio-Rad Gel Doc imager.

### Retron Acquisition Experiments

Cells were transformed sequentially: first with the RT expression plasmid (pSLS.400 or pSLS.402), and second with the ncRNA and Cas1+2 expression plasmid (eg. pSLS.416). For the -RT condition, cells were only transformed with an ncRNA and Cas1+2 expression plasmid (e.g. pSLS.416). For testing acquisition of retron-derived spacers in figures 1e-f, cells with RT, ncRNA, and Cas1+2 expression plasmids were grown overnight (16 hours) in 3 mL LB with antibiotics and inducers IPTG and arabinose, from individual clones on plates. In the morning, 240 uL of overnight culture was diluted into 3 mL fresh media with antibiotics, IPTG, and arabinose and grown for 2 hours. After 2 hours, 320 uL of culture was diluted into 3 mL fresh media with antibiotics and erythromycin (no erythromycin was used in the -RT condition) and grown for 8 hours. After 8 hours, culture was diluted 1:1000 into 3 mL LB with antibiotics and without inducers and grown overnight (16 hours). In the morning, 25 uL of culture was mixed with 25 uL of water, heated to 95C for 5 minutes to lyse cells, cooled, and frozen at -20C for later analysis. For data presented in Figures 2b-d and 3i, cells were grown overnight (16 hours) in 3 mL LB with antibiotics and inducers IPTG and arabinose, from individual clones on plates. In the morning, 240 uL of overnight culture was diluted into 3 mL fresh media with antibiotics, IPTG, and arabinose and grown for 2 hours. After 2 hours, 320 uL of culture was diluted into 3 mL fresh media with antibiotics and erythromycin and grown for 2 (rather than 8) hours. At this point, 25 uL of culture was mixed with 25 uL of water, heated to 95C for 5 minutes to lyse cells, cooled, and frozen at -20C for later analysis.

For the 24-hour time course experiment, the experiment was broken into two halves: the first 9 hours, and the final 15 hours. For the entirety of the time course, cells were grown in media with antibiotics and inducers (arabinose, IPTG, and erythromycin). For the first 9-hour samples, cultures were grown starting from single colonies added to 0.5 mL of media. These cultures were sampled every 1.5 hours until hour 9, with 1 mL of media added at hour 3 and 1.5 mL of media added at hour 6. For the final 15-hour samples, 3 mL of media was inoculated with single colonies from plates and grown for 9 hours. Starting at hour 9, samples were taken every 1.5 hours until hour 24. At hour 16.5, 200 uL of culture was diluted into 1.5 mL of fresh media and the experiment continued in the new tube. At hour 21, 1 mL media was added to the culture.

### Oligo Prespacer Feeding

For spacer acquisition experiments using exogeneous DNA prespacers (purified RT-DNA or synthetic oligos), cells containing pWUR1+2 were grown overnight from individual colonies on plates. In the morning, 100 uL of overnight culture was diluted into 3 mL LB with antibiotics, IPTG, and arabinose. Cells were grown with inducers for 2 hours. For each electroporation, 1 mL of culture was pelleted and resuspended in water. Cells were washed a second time by pelleting and resuspension, then pelleted one final time and resuspended in 50 uL of prespacer DNA solution at a concentration of 6.25 uM of single-stranded RT-DNA. All wash steps were done using ice cold water, all centrifugation steps were done in a centrifuge chilled to 4C, and samples kept on ice until electroporation was complete. The cell-DNA mixture was transferred to a 1 mm gap cuvette (Bio-Rad 1652089) and electroporated using a Bio-Rad gene pulser set to 1.8 kV and 25 uF with pulse controller at 200 Ohms. After electroporation, cells were recovered in 3 mL of LB without antibiotics for 2 hours. Then, 25 uL of culture was mixed with 25 uL of water, heated to 95C for 5 minutes to lyse cells, cooled, and frozen at -20C for later analysis.

### Signal Promoter Strength Measurement

bSLS.114 was transformed with Marionette plasmid^34^ (pAJM.683, pAJM.011, or pAJM.771) and grown overnight in LB with antibiotic (kanamycin). In the morning, 60ul of overnight culture was added to 2 tubes of 3 ml LB, one with antibiotic and inducer and the other with antibiotic and no inducer. The cultures were grown for 2 hours, 1ml of cell suspension pelleted (8000g for 1 min) and resuspended twice in 1mL PBS, and OD600 (600nm absorbance) and YFP fluorescence (513nm excitation/538nm emission) measurements were taken on a Molecular Devices SpectraMax i3 plate reader using a black, clear bottom 96-well plate and 200ul of resuspended cells. OD600 was measured using a kinetic scan for 2 minutes, taking measurements every 25 seconds, with a 1 second shake in between. Fluorescence was measured as a kinetic scan for 2 minutes, taking measurements every 20 seconds, with a 1 second shake in between.

### Recording System Fitness Measurements

Cells were transformed with plasmids (sequentially in the case of multiple plasmids). Single colonies were picked from plates and grown overnight in 3 mL of LB with antibiotics and without inducers. In the morning, 60 uL of culture was added to 3 mL of LB with antibiotics and left to sit at room temp for ∼4 hours. Then, cultures were transferred to a shaking incubator and grown for 2 hours. After 2 hours, the OD600 of each culture was measured using a NanoDrop-2000c spectrophotometer and cultures diluted to an OD600 of 0.05 in LB with antibiotics. Next, inducers were added at the appropriate strength and 200 uL of each culture (OD600 = 0.05 with antibiotics and inducers) was transferred to a clear-bottomed 96-well plate. The 96-well plate was loaded onto a Molecular Devices SpectraMax i3 plate reader set at 37C, and OD600 measurements taken every 2.5 minutes for the next 15 hours, with a 30 second shake before each reading.

### Temporal Recordings

Cells were transformed sequentially, first with pSBK.134 or pSBK.136 and then with pSBK.079. For recording, single colonies were picked from plates and grown overnight in 3 mL of LB with antibiotics and without inducers. In the morning, 150 uL of culture was diluted into 3 mL of LB with antibiotics and appropriate inducers (Fig. 4) and grown for 8 hours. After 8 hours, 60 uL of culture was diluted into 3 mL of LB with appropriate inducers and grown overnight (16 hours). In the morning, 150 uL of culture was diluted into 3 mL of LB with appropriate inducers (for second day of expression) and grown for 8 hours. Samples were collected at this 24-hour timepoint. 25 uL of culture was mixed with 25 uL of water, heated to 95C for 5 minutes to lyse cells, cooled, and frozen at -20C for later analysis. After 8 hours, 60 uL of culture was diluted into 3 mL of LB with appropriate inducers and grown overnight (16 hours). In the morning, 25 uL of culture was mixed with 25 uL of water, heated to 95C for 5 minutes to lyse cells, cooled, and frozen at -20C for later analysis.

### Computational Model of Retron Recording

A computational model of retron recording was written using Python 3 and the following modules, packages, and libraries: numpy, matplotlib, random, itertools, and xlsxwriter. For modeling, we assume that spacer acquisition is well approximated as a Poisson process in which acquisitions occur at some average rate over time, but where the precise timing of these events is random and independent of the timing of previous events. We believe this is a fair approximation of spacer acquisition due to the overall low rate of spacer acquisition in the retron recording system (single digit percentages of arrays expanded over 24 hours), the demonstrated ability of CRISPR arrays to be multiply expanded, and the current understanding of CRISPR adaptation indicating that acquisitions occur one at a time. To simulate spacer acquisition in a population of cells, we first define a “Cell” class of which each instance possesses an attribute called an “array”. The user defines the following parameters of the recording experiment: number of cells, rates of acquisition (in units of integrations per hour per cell) of retron-derived signal “A” with and without inducer present, rates of acquisition of retron-derived signal “B” with and without inducer present, rate of acquisition of non-retron-derived signal “N”, time of induction of signal “A”, time of induction of signal “B”, and order of induction (e.g. “A” before “B”). For the first epoch, each “Cell” instance samples three different Poisson distributions (one each for signals “A”, “B”, and “N”) to determine the number of spacers of each type which are added to its “array” during the epoch. The order of these spacers is then randomized and appended to the “array”. For example: when the order of induction is “A” before “B”, the cell is subject to the following rates of acquisition: “A” with inducer, “B” without inducer, and “N”. For each signal (“A”, “B”, and “N”) the cell samples a Poisson distribution defined by the probability mass function 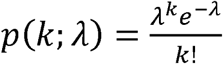 where *K* is the number of acquisitions of that signal (*K*= 0,1, 2 …) and *⁁* is the expected number of acquisitions of that signal (defined as the rate of acquisition of a given signal times the length of the epoch). It is fair to randomize the order of acquisitions occurring in each epoch, prior to appending them to the array, because the timing of the events is random by definition. For example: given that a cell acquires one “A” spacer and one “N” spacer in an interval with constant rates of acquisition of “A” and “N”, it is equally likely that “A” comes before “N” as it is that “N” comes before “A”. After acquisitions during the first epoch are completed, the process is repeated for the second epoch (using the relevant rates of acquisition for all three signals). At this point, the arrays are complete and ready for analysis using the ordering rules. Recordings, replicates, and ordering rule analysis were simulated using purpose-built scripts to investigate parameters of interest. Relevant data was exported to Excel sheets for further analysis and visualized using GraphPad Prism.

### Long-Term Passage for Data Stability

24+24-hour, “A”-then-”B” recordings were made in bSLS.114 cells harboring plasmids pSBK.134 and pSBK.079 as described previously. At hour 48, 25 uL of culture was mixed with 25 uL of water, heated to 95C for 5 minutes to lyse cells, cooled, and frozen at -20C for later analysis. 500 uL of culture was combined with 500 uL of 50% glycerol and frozen at -80C for future outgrowth. To begin long-term culture, recording glycerol stocks stored at -80C were thawed at room temperature, 100 ul of thawed cells added to 25 mL of LB with antibiotics, and the culture left to shake at 37C for 24 hours. Every day for the next 14 days, 25 ul of culture was sampled, boiled, and frozen as above, and 50 uL of culture added to 25 mL of fresh LB with antibiotics (ratio of 1:500, yielding roughly 9 generations per day). Samples from days 0, 2, 5, and 9 were sequenced and analyzed.

### Analysis of Spacer Acquisition

Analysis of spacer acquisition was conducted by sequencing a library of all CRISPR arrays in an experimental population using an Illumina MiSeq instrument. Libraries were created by amplifying a region of the genomic CRISPR array using PCR, then indexed using custom indexing oligos. Up to 192 conditions were run per flow cell. A list of oligo prespacers and primers can be found in Supplementary Table 4.

### Processing and Analysis of MiSeq Data

Sequences were analyzed using custom Python software, which will be available on GitHub upon peer-reviewed publication. In brief, newly acquired spacer sequences were extracted from array sequences based on their position between identifiable repeats and compared to preexisting spacers in the array. In this preliminary analysis, metrics were collected including number of expansions in arrays (unexpanded, single, double, and triple expanded) and proportion of each present in the library. Sequenced arrays were sorted into subcategories based on these characteristics (e.g. doubly expanded with first three repeats identifiable) for further analysis. Next, to determine number of retron-derived spacers and the order of spacers in multiply expanded arrays, two different analyses were used: one strict and one lenient. In the strict analysis (used in figures 1, 2, and 3) a retron-derived spacer is defined to be a spacer which contains the 23-base core region of the hypothetical prespacer structure from a given retron (with three mismatches or indels allowed). In the lenient analysis (used in figures 4 and 5) a retron-derived spacer is defined to be a spacer which contains an 11-base region of the hypothetical prespacer consisting of the 7-base barcode region and 2 bases on either side (with one mismatch or indel allowed). The order of spacers in multiply expanded arrays is then reported (e.g. Leader-NNA) and these data are used to complete the ordering rule analysis.

## Data Availability

All data supporting the findings of this study are available within the article and its supplementary information, or will be made available from the authors upon request. Sequencing data associated with this study are available in the NCBI SRA (PRJNA838025).

## Code Availability

Custom code to process and analyze data from this study is available on GitHub (https://github.com/Shipman-Lab/Spacer-Seq).

## ACKNOWLEDGEMENTS

Work was supported by funding from the Simons Foundation Autism Research Initiative (SFARI) Bridge to Independence Award Program, the Pew Biomedical Scholars Program, the NIH/NIGMS (1DP2GM140917-01), and the UCSF Program for Breakthrough Biomedical Research. S.L.S. is a Chan Zuckerberg Biohub investigator and acknowledges additional funding support from the L.K. Whittier Foundation. S.K.L. was supported by an NSF Graduate Research Fellowship (2034836). S.C.L was supported by a Berkeley Fellowship for Graduate Study. We thank Kathryn Claiborn for editorial assistance.

## AUTHOR CONTRIBUTIONS

S.L.S. conceived the study with J.N. and G.M.C. contributing. S.B.K. and S.L.S designed experiments and analyzed all data. Contributions to data collection were made from S.K.L. (Extended Data Figure 4), C.B.F. (Extended Data Figure 5), and S.L.S. (Figure 1c, e; Figure 3b-e). S.B.K. collected all other data. E.L., M.G.S., and S.C.L. performed preliminary experiments not included in the figures. S.B.K. performed the computational modeling of recordings. S.B.K. wrote the manuscript with input from all co-authors.

## Extended Data Figure Legends

**Extended Data Figure 1. Accompaniment to Figure 1. a**. Hypothetical Eco1 wild-type ncRNA-linked RT-DNA structure. **b**. Hypothetical Eco1 v32 ncRNA-linked RT-DNA structure and hypothetical duplexed RT-DNA prespacer structure. Nucleotides that are altered from wild-type Eco1 are shown in orange. **c**. Hypothetical Eco1 v35 ncRNA-linked RT-DNA structure and hypothetical duplexed RT-DNA prespacer structure. Nucleotides that are altered from wild-type Eco1 are shown in green.

**Extended Data Figure 2. Accompaniment to Figure 2. a**. Hypothetical barcoded Eco1 v35 ncRNA-linked RT-DNA structure and hypothetical duplexed RT-DNA prespacer structure. Bases used to barcode retrons are shown in red.

**Extended Data Figure 3. Accompaniment to Figure 3.** a. Hypothetical wild-type Eco4 ncRNA-linked RT-DNA structure. ExoVII-dependent RT-DNA cleavage site is shown as a red slash. **b**. Eco4-derived spacer sequences and orientations. Bases are colored to match Figure 3f. **c**. Proportion of Eco4-derived spacers in each orientation. Open circles are individual biological replicates.

**Extended Data Figure 4. Change in YFP fluorescence when expressed using inducible promoters**. The Y-axis shows fluorescence (in arbitrary units) normalized to culture density (OD600).

**Extended Data Figure 5. Growth curves (upper plot) and max growth rates (lower plot) of E. coli with different combinations of retron recording components and inducers**. In growth curve plots the solid line is the mean OD600 of 3 biological replicates, with dotted lines showing the standard deviation. In maximum growth rate plots, each symbol is a single biological replicate. Bars show the mean and standard deviation. Statistically significant differences in maximum growth rate, as calculated by Tukey’s multiple comparison’s test, are highlighted. **a**. Growth kinetics of E. coli with different combinations of retron recording plasmids, all without inducers. **b**. Growth kinetics of E. coli with recording plasmid pSBK.079, with and without inducers. **c**. Growth kinetics of E. coli with signal plasmid pSBK.134, with and without inducers. Only one biological replicate is present in condition “pSBK.134 + aTc” (pink). **d**. Growth kinetics of E. coli with signal plasmid pSBK.136, with and without inducers. **e**. Growth kinetics of E. coli with signal plasmid pSBK.134 and recording plasmid pSBK.079, with and without inducers. **f**. Growth kinetics of E. coli with signal plasmid pSBK.136 and recording plasmid pSBK.079, with and without inducers.

**Extended Data Figure 6. Accompaniment to Figure 4. a**. Ordering rules for pSBK.134 “A”-before-”B” replicates. The scores for each rule, and the composite score, are shown for each individual replicate. X-containing boxes indicate that no informative arrays, for that particular rule, were present in that replicate. **b**. As in panel (a), ordering rules for pSBK.134 “B”-before-”A” replicates. **c**. As in panel (a), ordering rules for pSBK.136 “A”-before-”B” replicates. **d**. As in panel (a), ordering rules for pSBK.136 “B”-before-”A” replicates.

**Extended Data Figure 7. Long-term stability of retron-derived recordings in CRISPR arrays. a**. Ordering rules for 24+24-hour, “A”-before-”B” recordings during post-recording multiday culture. Individual and composite scores are shown for samples taken on days 0, 2, 5, and 9 of culture. Each open circle represents the score, for that rule, from a single biological replicate. A total of 3 biological replicates are shown here. **b**. Changes in ordering rule scores over time in biological replicate 1. **c**. Changes in ordering rule scores over time in biological replicate 2. **d**. Changes in ordering rule scores over time in biological replicate 3.

