## Supplementary figures and images for "Recording gene expression order in DNA by CRISPR addition of retron barcodes"

### Extended Data Figure 1

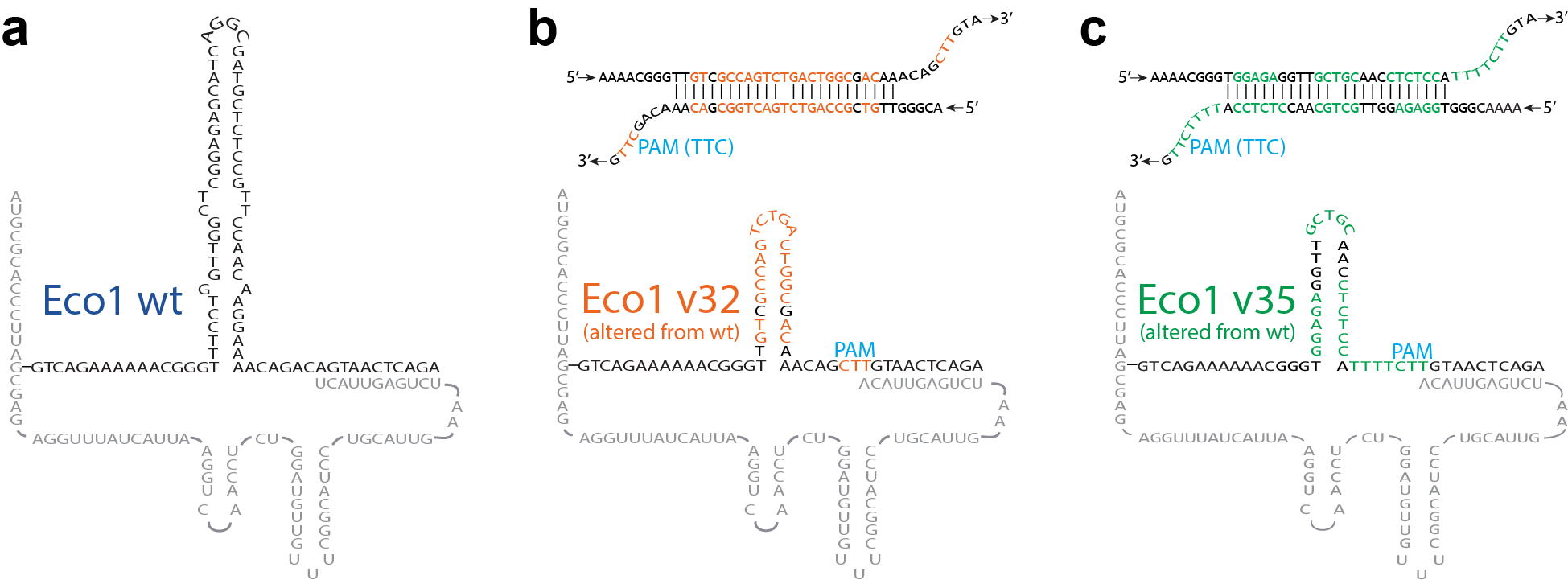

### Extended Data Figure 2

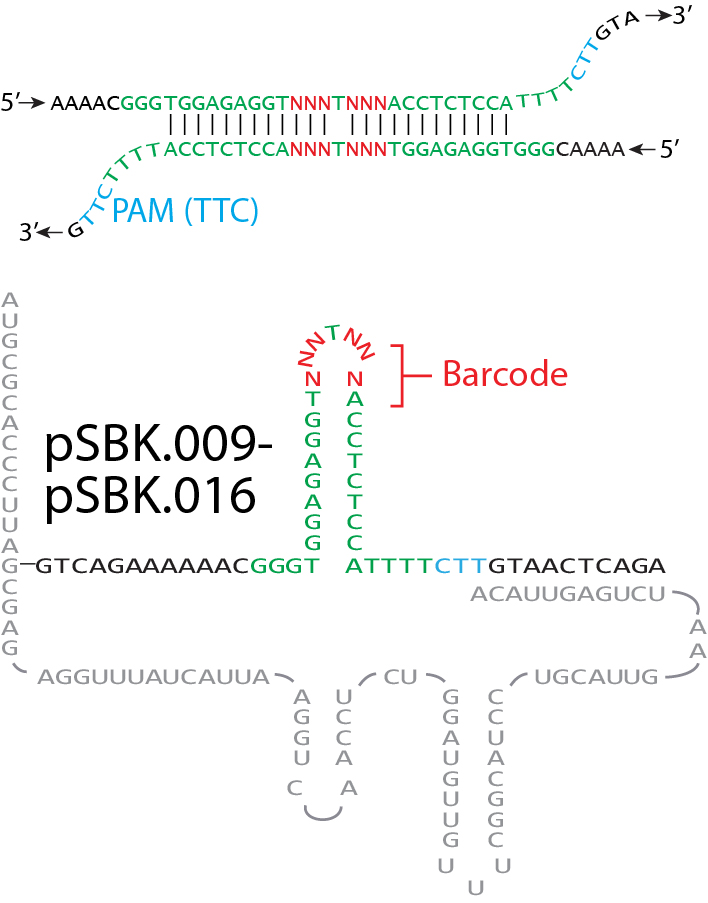

### Extended Data Figure 3

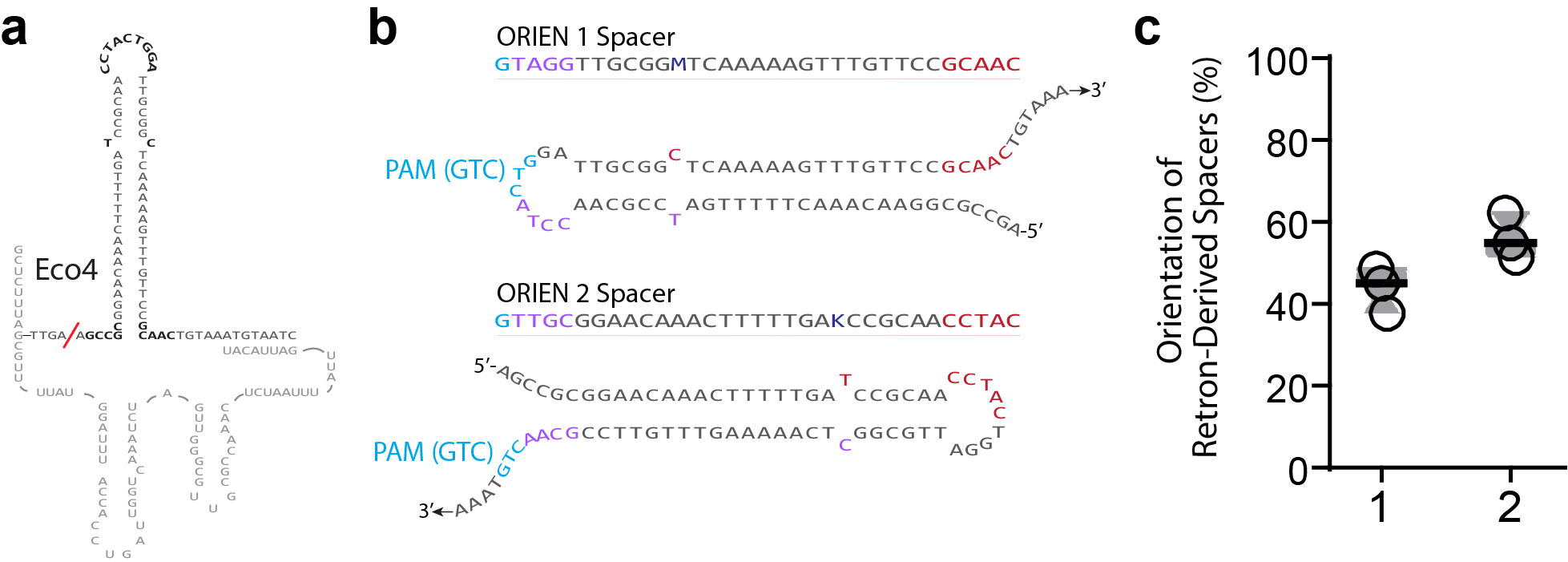

### Extended Data Figure 4

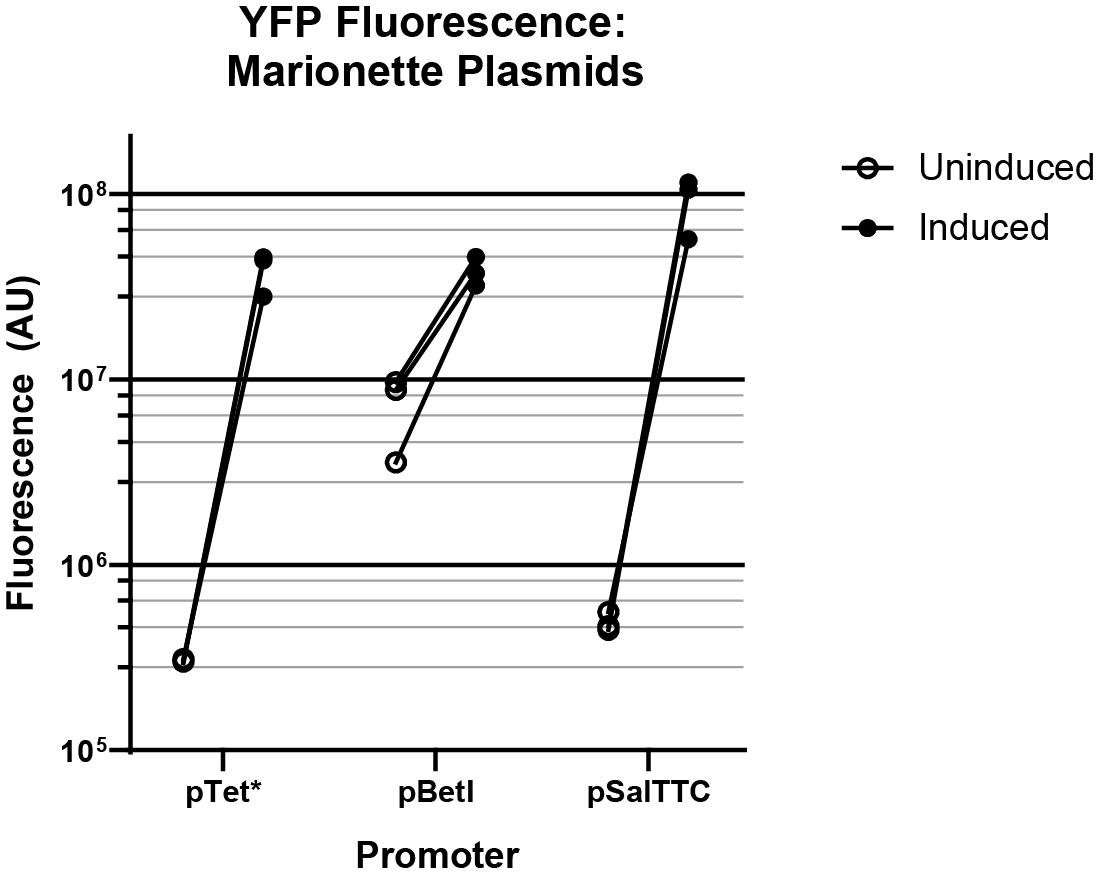

### Extended Data Figure 5

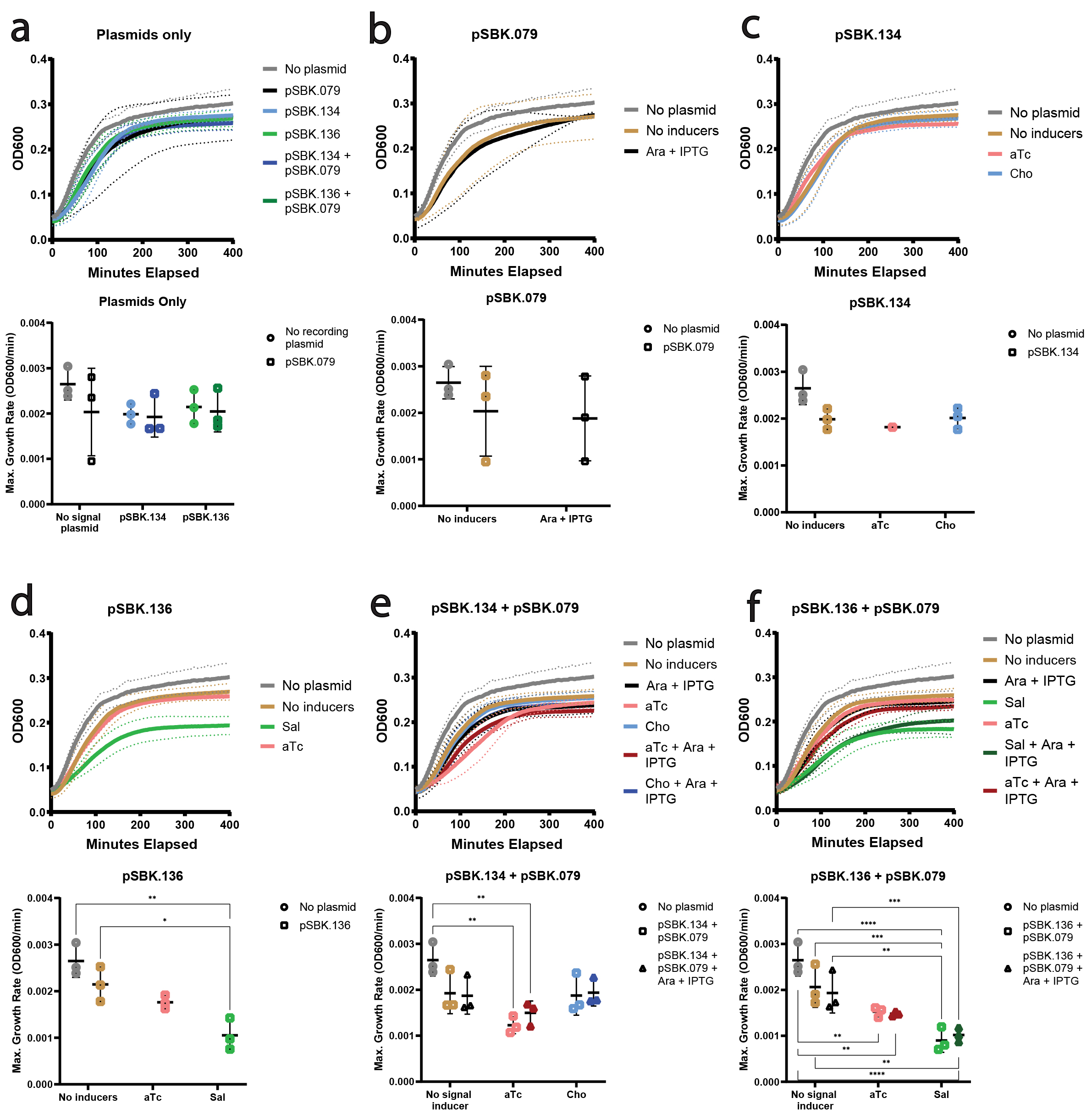

### Extended Data Figure 6

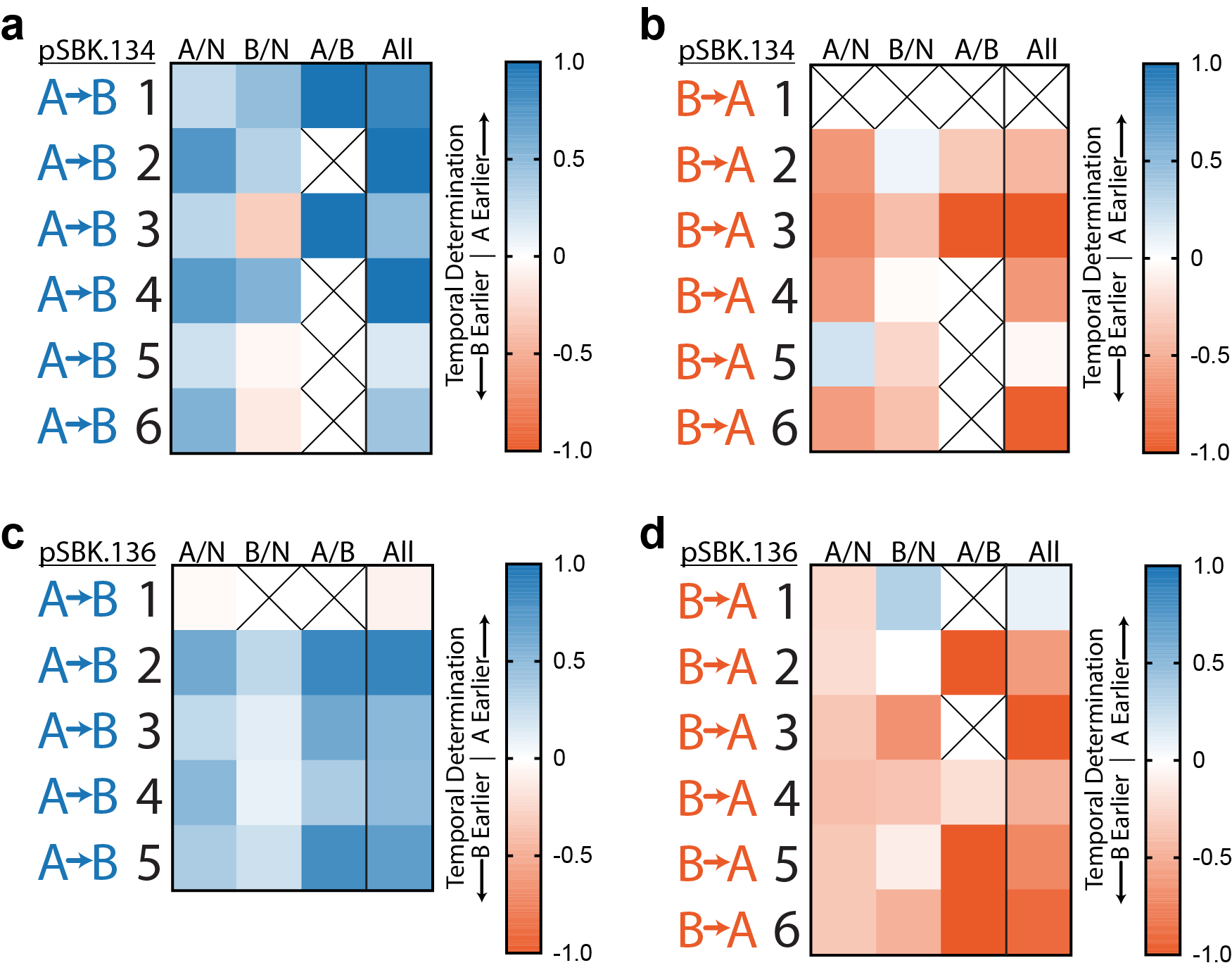

### Extended Data Figure 7

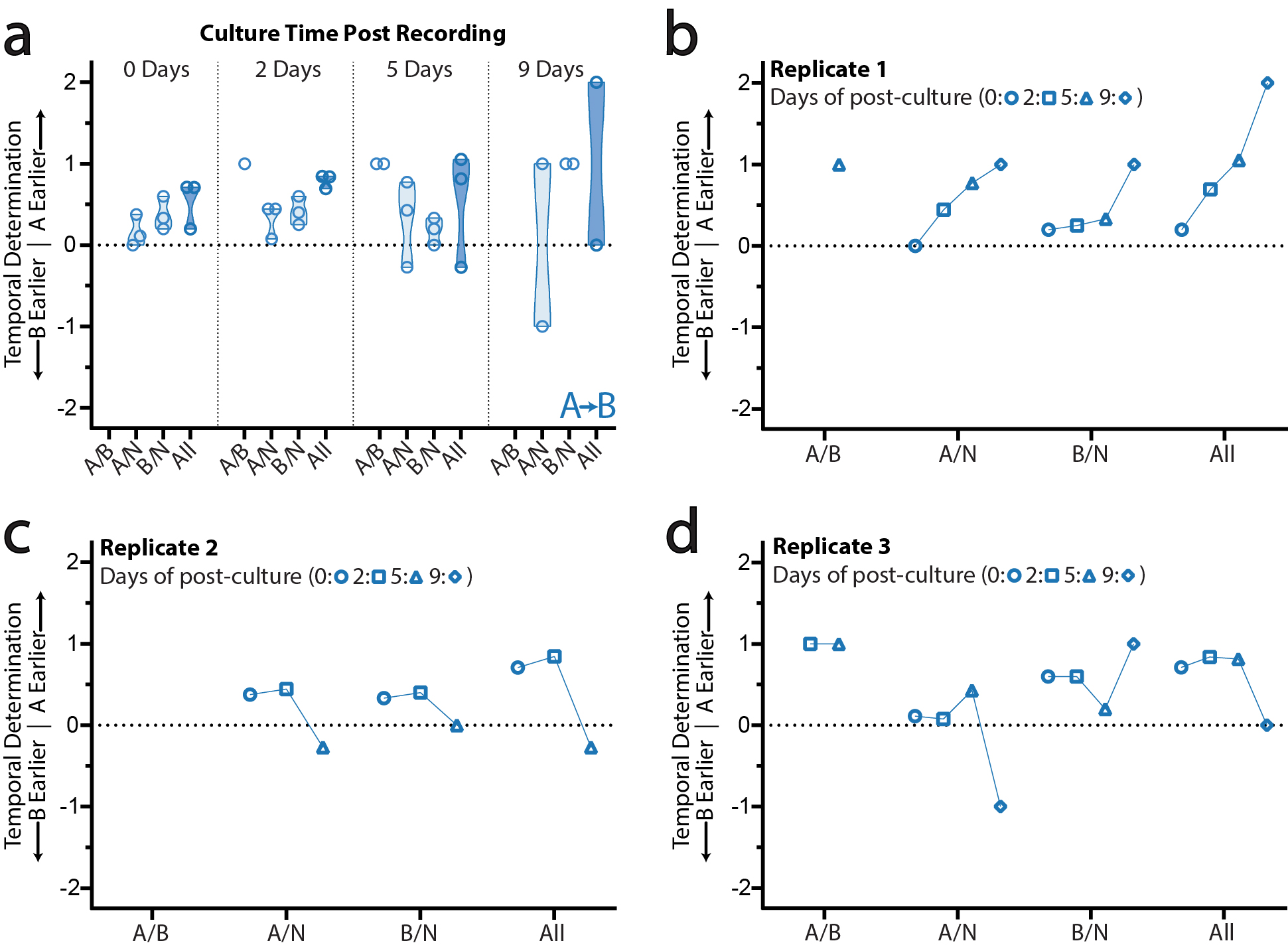
