## Supplementary Information for "Recording gene expression order in DNA by CRISPR addition of retron barcodes"

This PDF file includes:

Supplementary Figure 1, Uncropped gels

Supplementary Tables 1 to 4

**Supplemental Figure 1, Uncropped gels from main figures.**

Figure 1c

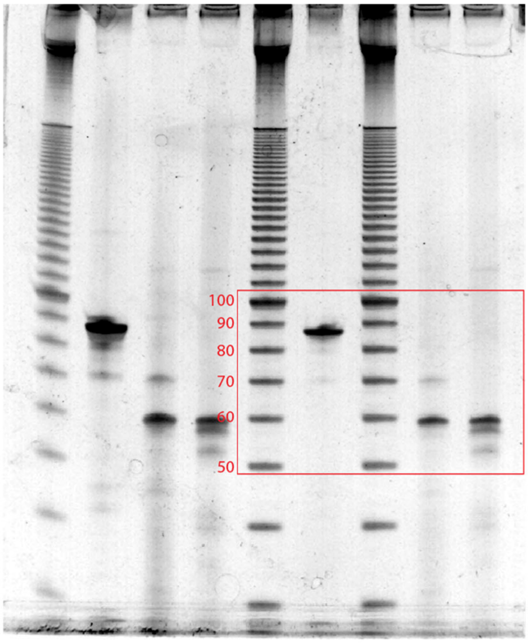


Figure 3c

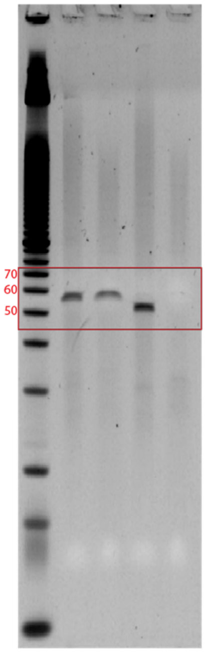


Figure 3h

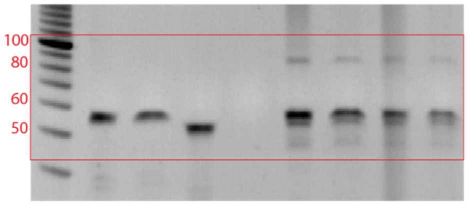

### Supplementary Table 1, Statistics

| | | Figure 1 | | | |
|---|---|---|---|---|---|
| Panel | Biological Replicates | Comparison | Test | *P* value | *P* value summary |
| e | 3 | v32 with RT versus without RT | unpaired t test | <0.0001 | **** |
| | 3 | v35 with RT versus without RT | unpaired t test | 0.0002 | *** |
| f | 5 | a1/a2 length: 12 versus 27 | unpaired t test | 0.0082 | ** |
| | | Figure 2 | | | |
| Panel | Biological Replicates | Comparison | Test | *P* value | *P* value summary |
| b | 3 | Non-retron derived spacers: effect of condition | one-way ANOVA | 0.5707 | ns |
| | | Retron derived spacers: effect of condition | one-way ANOVA | 0.0004 | *** |
| | | Follow-up: v35 "A" with RT versus without RT | Dunnett's multiple comparisons test (corrected) | <0.0001 | **** |
| | | Follow-up: v35 "A" versus "B" 1 | Dunnett's multiple comparisons test (corrected) | 0.029 | * |
| | | Follow-up: v35 "A" versus "B" 2 | Dunnett's multiple comparisons test (corrected) | 0.0229 | * |
| | | Follow-up: v35 "A" versus "B" 3 | Dunnett's multiple comparisons test (corrected) | 0.0119 | * |
| | | Follow-up: v35 "A" versus "B" 4 | Dunnett's multiple comparisons test (corrected) | 0.0357 | * |
| | | Follow-up: v35 "A" versus "B" 5 | Dunnett's multiple comparisons test (corrected) | 0.0287 | * |
| | | Follow-up: v35 "A" versus "B" 6 | Dunnett's multiple comparisons test (corrected) | 0.2986 | ns |
| | | Follow-up: v35 "A" versus "B" 7 | Dunnett's multiple comparisons test (corrected) | 0.086 | ns |
| | | Follow-up: v35 "A" versus "B" 8 | Dunnett's multiple comparisons test (corrected) | 0.0007 | *** |
| | | Figure 3 | | | |
| Panel | Biological Replicates | Comparison | Test | *P* value | *P* value summary |
| b | 4-5 | Effect of source (retron versus oligo) | two-way ANOVA | <0.0001 | **** |
| | | Effect of location (LP/M/LD) | two-way ANOVA | <0.0001 | **** |
| | | Follow-up: retron versus oligo, LP +/- ratio | Sidak's multiple comparisons test (corrected) | >0.9999 | ns |
| | | Follow-up: retron versus oligo, M +/- ratio | Sidak's multiple comparisons test (corrected) | 0.9985 | ns |
| | | Follow-up: retron versus oligo, M +/- ratio | Sidak's multiple comparisons test (corrected) | <0.0001 | **** |
| d | 2-5 comparison to oligo in panel b | Effect of source (retron versus oligo) | two-way ANOVA | 0.0439 | * |
| | | Effect of location (LP/M/LD) | two-way ANOVA | <0.0001 | **** |
| | | Follow-up: retron versus oligo, LP +/- ratio | Sidak's multiple comparisons test (corrected) | >0.9999 | ns |
| | | Follow-up: retron versus oligo, M +/- ratio | Sidak's multiple comparisons test (corrected) | >0.9999 | ns |
| | | Follow-up: retron versus oligo, M +/- ratio | Sidak's multiple comparisons test (corrected) | 0.005 | ** |
| e | 2-5 | Effect of condition | one-way ANOVA | 0.33 | ns |
| | | Follow-up: oligo versus retron -DBR1 | Dunnett's multiple comparisons test (corrected) | 0.2858 | ns |
| | | Follow-up: oligo versus retron +DBR1 | Dunnett's multiple comparisons test (corrected) | 0.5715 | ns |
| g | 3 | Effect of source (retron versus oligo) | two-way ANOVA | 0.0394 | * |
| | | Effect of location (LP/M/LD) | two-way ANOVA | 0.0083 | ** |
| | | Follow-up: retron versus oligo, LP +/- ratio | Sidak's multiple comparisons test (corrected) | >0.9999 | ns |
| | | Follow-up: retron versus oligo, M +/- ratio | Sidak's multiple comparisons test (corrected) | >0.9999 | ns |
| | | Follow-up: retron versus oligo, M +/- ratio | Sidak's multiple comparisons test (corrected) | 0.0053 | ** |
| i | 3 | Eco4 with RT versus without RT | unpaired t test | 0.0197 | * |
| | | Figure 4 | | | |
| Panel | Biological Replicates | Comparison | Test | *P* value | *P* value summary |
| b | 4 | "A" (aTc) versus "B" (Cho) | unpaired t test | 0.2704 | ns |
| c | 4 | Effect of Time on Retron-Derived Acquisition Rate (overall) | two-way ANOVA | 0.4172 | ns |
| | | Effect of Source on Retron-Derived Acquisition Rate (overall) | two-way ANOVA | 0.9312 | ns |
| | | Effect of Interaction Between Time and Source on Retron-Derived Acquisition Rate | two-way ANOVA | 0.0007 | *** |
| | | Follow-up: Rate of "A" Acquisitions, 24h versus 48h | Sidak's multiple comparisons test (corrected) | 0.005 | ** |
| | | Follow-up: Rate of "B" Acquisitions, 24h versus 48h | Sidak's multiple comparisons test (corrected) | 0.0445 | * |
| d | 4 | Effect of Time on Retron-Derived Acquisition Rate (overall) | two-way ANOVA | 0.0002 | *** |
| | | Effect of Source on Retron-Derived Acquisition Rate (overall) | two-way ANOVA | >0.9999 | ns |
| | | Effect of Interaction Between Time and Source on Retron-Derived Acquisition Rate | two-way ANOVA | <0.0001 | **** |
| | | Follow-up: Rate of "A" Acquisitions, 24h versus 48h | Sidak's multiple comparisons test (corrected) | <0.0001 | **** |
| | | Follow-up: Rate of "B" Acquisitions, 24h versus 48h | Sidak's multiple comparisons test (corrected) | 0.888 | ns |
| e | 4 | Effect of Time on Non-Retron-Derived Acquisition Rate (overall) | two-way ANOVA | 0.0633 | ns |
| | | Effect of Inducer Order on Non-Retron-Derived Acquisition Rate (overall) | two-way ANOVA | >0.9999 | ns |
| | | Effect of Interaction Between Time and Order on Non-Retron-Derived Acquisition Rate | two-way ANOVA | 0.5455 | ns |
| | | Follow-up: Rate of "N" Acquisitions A->B, 24h versus 48h | Sidak's multiple comparisons test (corrected) | 0.1602 | ns |
| | | Follow-up: Rate of "N" Acquisitions B->A, 24h versus 48h | Sidak's multiple comparisons test (corrected) | 0.5562 | ns |
| g | 4 | "A" (Sal) versus "B" (aTc) | unpaired t test | <0.0001 | **** |
| h | 3 | Effect of Time on Retron-Derived Acquisition Rate (overall) | two-way ANOVA | 0.0003 | *** |
| | | Effect of Source on Retron-Derived Acquisition Rate (overall) | two-way ANOVA | >0.9999 | ns |
| | | Effect of Interaction Between Time and Source on Retron-Derived Acquisition Rate | two-way ANOVA | 0.005 | ** |
| | | Follow-up: Rate of "A" Acquisitions, 24h versus 48h | Sidak's multiple comparisons test (corrected) | 0.3036 | ns |
| | | Follow-up: Rate of "B" Acquisitions, 24h versus 48h | Sidak's multiple comparisons test (corrected) | 0.0002 | *** |
| i | 4 | Effect of Time on Retron-Derived Acquisition Rate (overall) | two-way ANOVA | 0.0473 | * |
| | | Effect of Source on Retron-Derived Acquisition Rate (overall) | two-way ANOVA | 0.392 | ns |
| | | Effect of Interaction Between Time and Source on Retron-Derived Acquisition Rate | two-way ANOVA | 0.0002 | *** |
| | | Follow-up: Rate of "A" Acquisitions, 24h versus 48h | Sidak's multiple comparisons test (corrected) | 0.1003 | ns |
| | | Follow-up: Rate of "B" Acquisitions, 24h versus 48h | Sidak's multiple comparisons test (corrected) | 0.0004 | *** |
| j | 4 | Effect of Time on Non-Retron-Derived Acquisition Rate (overall) | two-way ANOVA | 0.1855 | ns |
| | | Effect of Inducer Order on Non-Retron-Derived Acquisition Rate (overall) | two-way ANOVA | 0.5823 | ns |
| | | Effect of Interaction Between Time and Order on Non-Retron-Derived Acquisition Rate | two-way ANOVA | 0.0007 | *** |
| | | Follow-up: Rate of "N" Acquisitions A->B, 24h versus 48h | Sidak's multiple comparisons test (corrected) | 0.09 | ns |
| | | Follow-up: Rate of "N" Acquisitions B->A, 24h versus 48h | Sidak's multiple comparisons test (corrected) | 0.0014 | ** |
| l | 6 | Effect of Order (A->B vs B->A) | two-way ANOVA | <0.0001 | **** |
| | | Effect of Rule | two-way ANOVA | 0.7179 | ns |
| | | Effect of Interaction | two-way ANOVA | 0.0044 | ** |
| | | Follow-up: A/N (A->B vs B->A) | Sidak's multiple comparisons test (corrected) | <0.0001 | **** |
| | | Follow-up: N/B (A->B vs B->A) | Sidak's multiple comparisons test (corrected) | 0.236 | ns |
| | | Follow-up: A/B (A->B vs B->A) | Sidak's multiple comparisons test (corrected) | <0.0001 | **** |
| | | Follow-up: All (A->B vs B->A) | Sidak's multiple comparisons test (corrected) | 0.0003 | *** |
| | | A->B Overall Values Different Than 0 | one-sample t test | 0.0065 | ** |
| | | B->A Overall Values Different Than 0 | one-sample t test | 0.0363 | * |
| m | 5 | Effect of Order (A->B vs B->A) | two-way ANOVA | <0.0001 | **** |
| | | Effect of Rule | two-way ANOVA | 0.9252 | ns |
| | | Effect of Interaction | two-way ANOVA | 0.0017 | ** |
| | | Follow-up: A/N (A->B vs B->A) | Sidak's multiple comparisons test (corrected) | 0.0011 | ** |
| | | Follow-up: N/B (A->B vs B->A) | Sidak's multiple comparisons test (corrected) | 0.1054 | ns |
| | | Follow-up: A/B (A->B vs B->A) | Sidak's multiple comparisons test (corrected) | <0.0001 | **** |
| | | Follow-up: All (A->B vs B->A) | Sidak's multiple comparisons test (corrected) | <0.0001 | **** |
| | | A->B Overall Values Different Than 0 | one-sample t test | 0.0327 | * |
| | | B->A Overall Values Different Than 0 | one-sample t test | 0.0116 | * |

#### Supplementary Table 2, Strains

| Name | Species | Parental Line | Genotype | Method |
|---|---|---|---|---|
| NEB 5-alpha | *E. coli* | DH5α | *fhuA2 Δ(argF-lacZ)U169 phoA glnV44 Φ80 Δ(lacZ)M15 gyrA96 recA1 relA1 endA1 thi-1 hsdR17* | |
| BL21-AI | *E. coli* | BL21 | *F-ompT hsdSB (rB- mB-) gal dcm araB::T7RNAP-tetA* | |
| bSLS.114 | *E. coli* | BL21-AI | *E. coli B F– ompT gal dcm lon hsdSB*($rB^-$ $mB^-$) [$malB^+$]$_{K\text{-}12}$ ($\lambda^S$) *araB::T7RNAP-tetA ΔEco1* | lambda Red recombinase mediated insertion of chloramphenicol resistance, marker excision by FLP |
| bMS.346 | *E. coli* | MG1655 | *MG1655 galKL187TAAL188TGA Δexol ΔrecJ araB^:pBAD>T7RNAP;Tet* | lambda Red recombination and phage P1 transduction |

#### Supplementary Table 3, Plasmids

| Name | Genes | Promoter | Inducer (working concentration) | Used in (panels) |
|---|---|---|---|---|
| pWUR 1+2 | Cas1 + Cas2 (Yosef et al. 2012) | T7/lac | L-arabinose (2 mg/mL), IPTG (1 mM) | 3g-i |
| pSLS.400 | Eco4 RT | pMphR | Erythromycin (400 μM) | 3g-i |
| pSLS.402 | Eco1 RT | pMphR | Erythromycin (400 μM) | 1c, 1e-i, 2b-d, 3b-e, 3h |
| pSLS.405 | 1: Eco1 ncRNA + Eco1 RT; 2: Cas1 + Cas2 | 1: T7/lac; 2: T7/lac | L-arabinose (2 mg/mL), IPTG (1 mM) | 1c, 3c, 3h |
| pSLS.407 | 1: Eco1 ncRNA v32; 2: Cas1 + Cas2 | 1: T7/lac; 2: T7/lac | L-arabinose (2 mg/mL), IPTG (1 mM) | 1c, 1e |
| pSLS.408 | 1: Eco1 ncRNA v35; 2: Cas1 + Cas2 | 1: T7/lac; 2: T7/lac | L-arabinose (2 mg/mL), IPTG (1 mM) | 1c, 1e-f |
| pSLS.416 | 1: Eco1 ncRNA v35, long a1/a2; 2: Cas1 + Cas2 | 1: T7/lac; 2: T7/lac | L-arabinose (2 mg/mL), IPTG (1 mM) | 1f-i, 2b-d, 3b, 3d-e |
| pSLS.419 | 1: Eco4 ncRNA; 2: Cas1 + Cas2 | 1: T7/lac; 2: T7/lac | L-arabinose (2 mg/mL), IPTG (1 mM) | 3g-i |
| pSBK.009 | 1: Eco1 ncRNA v35, long a1/a2, barcode 1 (CCT-AGG); 2: Cas1 + Cas2 | 1: T7/lac; 2: T7/lac | L-arabinose (2 mg/mL), IPTG (1 mM) | 2b-d |
| pSBK.010 | 1: Eco1 ncRNA v35, long a1/a2, barcode 2 (GCT-AGC); 2: Cas1 + Cas2 | 1: T7/lac; 2: T7/lac | L-arabinose (2 mg/mL), IPTG (1 mM) | 2b-d |
| pSBK.011 | 1: Eco1 ncRNA v35, long a1/a2, barcode 3 (CTG-CAG); 2: Cas1 + Cas2 | 1: T7/lac; 2: T7/lac | L-arabinose (2 mg/mL), IPTG (1 mM) | 2b-d |
| pSBK.012 | 1: Eco1 ncRNA v35, long a1/a2, barcode 4 (GTG-CAC); 2: Cas1 + Cas2 | 1: T7/lac; 2: T7/lac | L-arabinose (2 mg/mL), IPTG (1 mM) | 2b-d |
| pSBK.013 | 1: Eco1 ncRNA v35, long a1/a2, barcode 5 (ACG-CGT); 2: Cas1 + Cas2 | 1: T7/lac; 2: T7/lac | L-arabinose (2 mg/mL), IPTG (1 mM) | 2b-d |
| pSBK.014 | 1: Eco1 ncRNA v35, long a1/a2, barcode 6 (CAG-TAG); 2: Cas1 + Cas2 | 1: T7/lac; 2: T7/lac | L-arabinose (2 mg/mL), IPTG (1 mM) | 2b-d |
| pSBK.015 | 1: Eco1 ncRNA v35, long a1/a2, barcode 7 (GAG-CTC); 2: Cas1 + Cas2 | 1: T7/lac; 2: T7/lac | L-arabinose (2 mg/mL), IPTG (1 mM) | 2b-d |
| pSBK.016 | 1: Eco1 ncRNA v35, long a1/a2, , barcode 8 (GCT-TGC); 2: Cas1 + Cas2 | 1: T7/lac; 2: T7/lac | L-arabinose (2 mg/mL), IPTG (1 mM) | 2b-d |
| pSBK.079 | 1: Eco1 RT; 2: Cas1 + Cas2 | 1: J23115; 2: T7/lac | L-arabinose (2 mg/mL), IPTG (1 mM) | 4b-e, 4g-j, 4l-m |
| pSBK.134 | A: Eco1 ncRNA v35, long a1/a2; B:Eco1 ncRNA v35, long a1/a2, barcode 6 | A: pTet*; B: pBetI | A: anhydrotetracycline (100 ng/mL); B: choline chloride (100 μM) | 4b-e, 4l |
| pSBK.136 | A: Eco1 ncRNA v35, long a1/a2; B:Eco1 ncRNA v35, long a1/a2, barcode 6 | A: pSalTTC; B: pTet* | A: sodium salicylate (1 mM); B: anhydrotetracycline (100 ng/mL) | 4g-j, 4m |

#### Supplementary Table 4, Primers

| Name | Sequence | Purpose |
|---|---|---|
| SPCR_MiSeq3_fow1 | CTTTCCCTACACGACGCTCTTCCGATCTNCATTAATTAATAATAGGTTATGTTTAGAGTGTTCC | Forward primer with Illumina adapter (#1 of set of 5) used to amplify BL21-AI CRISPR array for sequencing |
| SPCR_MiSeq3_fow2 | CTTTCCCTACACGACGCTCTTCCGATCTNNCATTAATTAATAATAGGTTATGTTTAGAGTGTTCC | Forward primer with Illumina adapter (#2 of set of 5) used to amplify BL21-AI CRISPR array for sequencing |
| SPCR_MiSeq3_fow3 | CTTTCCCTACACGACGCTCTTCCGATCTNNNCATTAATTAATAATAGGTTATGTTTAGAGTGTTCC | Forward primer with Illumina adapter (#3 of set of 5) used to amplify BL21-AI CRISPR array for sequencing |
| SPCR_MiSeq3_fow4 | CTTTCCCTACACGACGCTCTTCCGATCTNNNNCATTAATTAATAATAGGTTATGTTTAGAGTGTTCC | Forward primer with Illumina adapter (#4 of set of 5) used to amplify BL21-AI CRISPR array for sequencing |
| SPCR_MiSeq3_fow5 | CTTTCCCTACACGACGCTCTTCCGATCTNNNNNCATTAATTAATAATAGGTTATGTTTAGAGTGTTCC | Forward primer with Illumina adapter (#5 of set of 5) used to amplify BL21-AI CRISPR array for sequencing |
| SPCR_MiSeq3_rev | GGAGTTCAGACGTGTGCTCTTCCGATCTGTGTCAACAATCGTTCCCTGATTGTC | Reverse primer with Illumina adapter used to amplify BL21-AI CRISPR array for sequencing |
| Eco1_v35_oligo | GTCAGAAAAAACGGGTGGAGAGGTTGCTGCAACCTCTCCATTTTCTTGTAACTCAGA | Test acquisition behavior of Eco1 v35 by electroporation |
| Eco4_wt_oligo | AGCCGCGGAACAAACTTTTTGATCCGCAACCTACTGGATTGCGGCTCAAAAAGTTTGTTCCGCAACTGTAAATGTAATC | Test acquisition behavior of Eco4 by electroporation |
